# Nature of active forces in tissues: how contractile cells can form extensile monolayers

**DOI:** 10.1101/2020.10.28.358663

**Authors:** Lakshmi Balasubramaniam, Amin Doostmohammadi, Thuan Beng Saw, Gautham Hari Narayana Sankara Narayana, Romain Mueller, Tien Dang, Minnah Thomas, Shafali Gupta, Surabhi Sonam, Alpha S. Yap, Yusuke Toyama, René-Marc Mège, Julia Yeomans, Benoît Ladoux

**Affiliations:** Institut Jacques Monod (IJM), CNRS UMR 7592 et Universite de Paris, 75013 Paris, France; Niels Bohr Institute, University of Copenhagen, Blegdamsvej 17, 2100 Copenhagen, Denmark; The Rudolf Peierls Centre for Theoretical Physics, University of Oxford, Parks Road, Oxford OX1 3PU, UK; Mechanobiology Institute (MBI), National University of Singapore, Singapore, 117411; National University of Singapore, Department of Biomedical Engineering, 4 Engineering Drive 3, Engineering Block 4, # 04-08, Singapore, 117583; Division of Cell and Developmental Biology, Institute for Molecular Bioscience, The University of Queensland, St. Lucia, Brisbane, QLD 4072, Australia; D Y Patil International University, Akurdi, Pune, India

## Abstract

Actomyosin machinery endows cells with contractility at a single cell level. However, at a tissue scale, cells can show either contractile or extensile behaviour based on the direction of pushing or pulling forces due to neighbour interactions or substrate interactions. Previous studies have shown that a monolayer of fibroblasts behaves as a contractile system^1^ while a monolayer of epithelial cells^2,3^ or neural crest cells behaves as an extensile system.^4^ How these two contradictory sources of force generation can coexist has remained unexplained. Through a combination of experiments using MDCK (Madin Darby Canine Kidney) cells, and in-silico modeling, we uncover the mechanism behind this switch in behaviour of epithelial cell monolayers from extensile to contractile as the weakening of intercellular contacts. We find that this switch in active behaviour also promotes the buildup of tension at the cell-substrate interface through an increase in actin stress fibers and higher traction forces. This in turn triggers a mechanotransductive response in vinculin translocation to focal adhesion sites and YAP (Yes-associated protein) transcription factor activation. Our studies also show that differences in extensility and contractility act to sort cells, thus determining a general mechanism for mechanobiological pattern formation during cell competition, morphogenesis and cancer progression.

## Main text

The ability of cell monolayers to self-organize, migrate and evolve depends crucially on the interplay between cell-matrix and cell-cell interactions^5–10^ which controls various phenomena including tissue morphogenesis,^11,12^ epithelial-mesenchymal transition,^5^ wound healing and tumor progression.^13^ Cells are active systems, engines that operate away from thermal equilibrium, transducing chemical energy into motion. Single isolated cells generate contractile force dipoles: the resultant of the forces due to actomyosin contraction, pulling on focal adhesion sites on the substrate, is typically a pair of approximately equal and opposite forces acting inwards along the cellular long axis^14^ (Figure 1a). It is reasonable to expect that contractile particles also generate contractile behaviour in the monolayer.^1^ However, at the collective cell level, epithelial monolayers display extensile behaviour^2^ i.e. the net force from the neighbours and substrate interaction act to elongate the cell further along its long axis (Figure 1b inset). This immediately poses the question of how such a crossover occurs as the emergence of such differences in active behaviour may be crucial in understanding biological processes such as tissue homeostasis, cell competition and self organization.^15^

**Figure 1.**
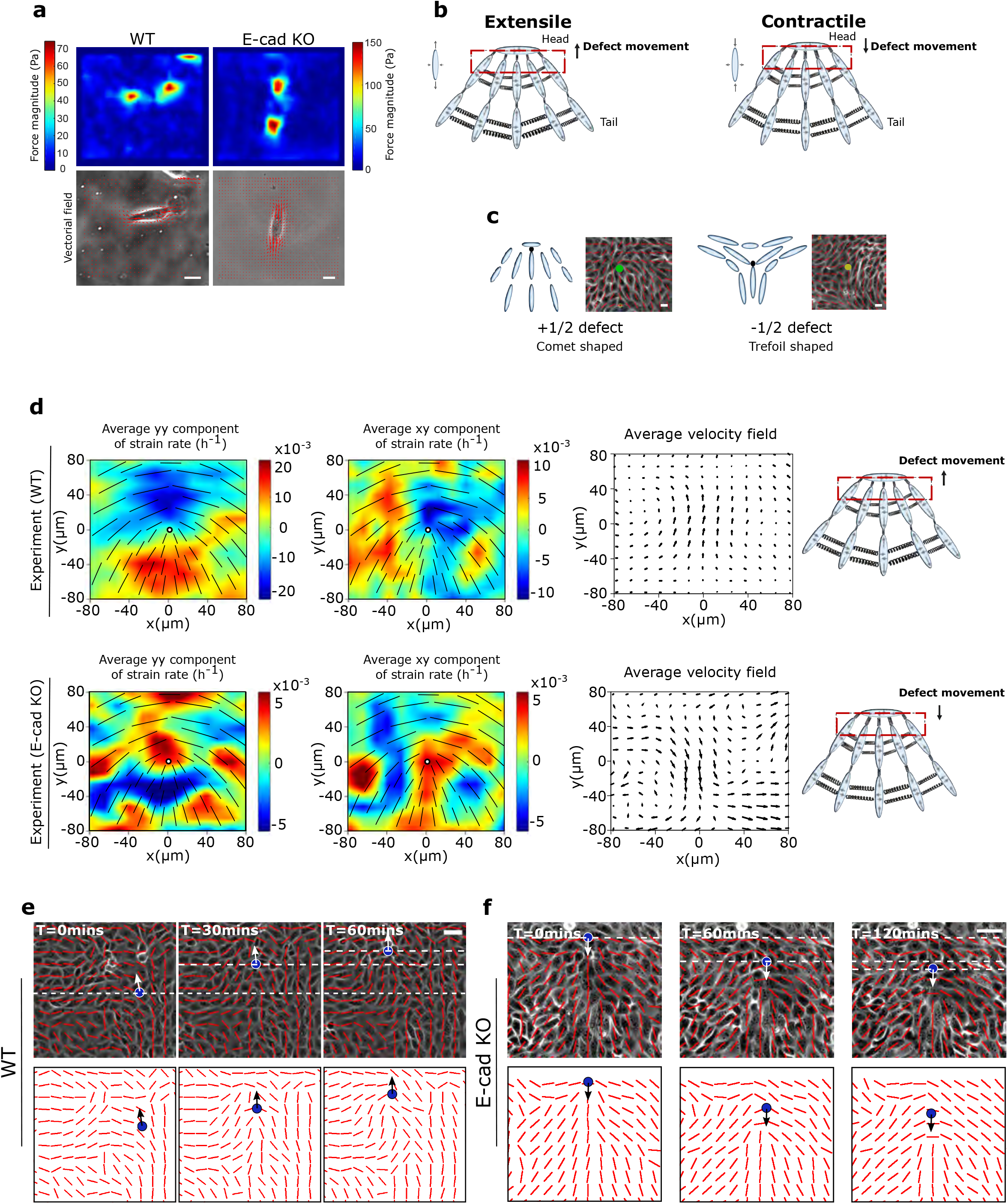
Active nematic behaviour of epithelial cellular systems changes from extensile to contractile in the absence of E-cadherin. a) Top, left and right: typical examples of traction force magnitude maps for a single MDCK WT and E-cadherin KO cell cultured on deformable PDMS surfaces. Bottom, left and right: vectorial maps of traction forces for a single MDCK WT and E-cadherin KO cell on a soft PDMS substrate. Scale bars, *20μm*. b) Schematic showing the defect movement based on force balance for an extensile active nematic system (left) and contractile active nematic system (right) with an inset of forces exerted on neighbours by an extensile (left) and contractile (right) nematic particle. c) Schematic (left) and experimental (right) images of *+1/2* defect (left, comet configuration) and *−1/2* defect (right, trefoil configuration). Scale bars, *20μm.* d) Average *yy*- and *xy*-components of strain rate map around *+ 1/2* defect obtained from experiments (left and middle respectively) and corresponding average flow field (right) for MDCK WT cells (top) (*n = 1934* defects from *2* independent experiments) and MDCK E-cadherin KO cells (bottom) (*n = 1,884* defects from *2* independent experiments). Schematic on the extreme right illustrates the movement of defects. Colour code is positive for stretching and negative for shrinkage. e, f) Experimental data for MDCK WT (e) and MDCK E-cadherin KO (f) monolayers. Top panels: phase contrast images of the cells overlaid with the average local orientation of the cells (red lines). Bottom panels: average local orientation of the cells (red lines). The blue circle shows the location of a *+1/2* defect and the corresponding arrow indicates the direction of motion of this defect over time. Dashed lines have been added for better reading of defect movement. Scale bars, *40μm*.

The extensility or contractility within cell populations are based on force balance as shown in Figure 1b and this can be determined by looking at the structure of flow fields around topological defects. Topological defects are singular points in the orientation field of the cell monolayers, where the orientation of cells were defined as the direction of their long axis (see Methods). Having identified the orientation of cells, we use the winding number parameter to identify the location of topological defects, using an automated defect detection method.^2^ In a cellular monolayer two types of topological defects predominate: comet-shaped defects and trefoils (Figure 1c), which correspond to topological defects in nematic liquid crystals with charges +1/2 and −1/2, respectively.^1−4,16^

Of relevance here in active systems, the active nature of cells results in a directed motion of the comet shaped defects. For extensile systems, the defects move in the direction of the head of the comet, while topological defects in contractile systems move towards the comet tail (Figure 1b). Thus, we measured the average flow field around the comet defects in Madin-Darby Canine Kidney Wild-Type (MDCK WT) monolayers using Particle Image Velocimetry (PIV). The flow and orientation field were obtained from time lapse imaging after they reached confluency and before the cells became isotropic as the monolayers grew too dense. The results show clearly that the comet-shaped defects move in the tail-to-head direction (Figure 1d,e, Supplementary Figure 1a, c, Video 1), indicating that at a collective level the MDCK monolayer behaves as an extensile active system despite forming contractile dipoles at a single cell level (Figure 1a). The extensile behaviour of comet-shaped defects has been recently reported for Human Bronchial Epithelial Cells (HBEC) as well,^3^ indicating it to be a property of epithelial monolayers. By contrast, the flow field around comet shaped defects in a monolayer of fibroblasts has an opposite flow direction - from head-to-tail of the comet - indicating that fibroblasts behave as a contractile system at the collective level (Supplementary Figure 2a), in agreement with previous studies.^1^ This difference in the direction of motion of defects is also reflected in the patterns of strain rates around the defects. While strain rate along the tail-to-head direction (*yy*-strain rates) show negative values at the head of a comet-shaped defect in MDCK WT monolayers indicating the presence of compression (Figure 1d), this is reversed for a monolayer of fibroblasts, where the *yy*-strain rate at the defect head is positive, indicating extensional deformation (Supplementary Figure 2a). But what causes epithelial cells to behave as an extensile system at the collective level, and mesenchymal cells as a contractile system, and what are the consequences during tissue organization are not well understood.

One fundamental difference between epithelial and mesenchymal cells is the ability of epithelial cells to form strong cell-cell adhesions through E-cadherin based junctional complexes, responsible for active intercellular force transmission.^17^ In order to discern the origin of extensile behaviour at a collective level, we performed laser ablation experiments on MDCK WT monolayers (Supplementary Figure 3a,b), where we observed higher recoil at shorter junctions in comparison to long junctions. These results highlight that the smaller cortical tension along long junctions gives rise to a tension distribution that leads to an extensile stress on the cell further elongating it. We therefore asked if weakening this intercellular adhesion in epithelial cells results in a (mesenchymal-like) contractile behaviour at the collective level. To test this, we inactivated the E-cadherin gene in MDCK cells using CRISPR-Cas9 which was validated through immunostaining and western blot analysis (Supplementary Figure 4a,b). MDCK E-cadherin Knock-Out (E-cad KO) cells can still maintain their contacts through another form of cadherin (cadherin 6),^18^ albeit with a significantly weaker adhesion strength as observed through the reduced level of *β*-catenin at adherens junctions (Supplementary Figure 4a,b), while still being able to form tight junctions (Supplementary Figure 4a). Strikingly, in these E-cad KO monolayers, the average flow field around comet defects switches direction compared to WT monolayers (Figure 1d and e, Supplementary Figure 1a,d, Video 2), indicating a contractile behaviour at the collective level similar to that of fibroblasts where the comet shaped defects move towards the tail direction (Supplementary Figure 2). This change in direction of the flow field around the defect was accompanied by changes in the average strain rate patterns which are positive (extensile deformation) around the head of a comet shaped defect in E-cadherin KO monolayers in comparison to WT monolayers where the strain rate is negative (compressive deformation) around the head of the defect (Figure 1d). Therefore, epithelial monolayers behave as an extensile system due to the presence of strong cell-cell adhesions and loosening this adhesion by removing E-cadherin results in a contractile behaviour.

In order to check that this switch from extensile to contractile behaviour is not only specific to MDCK cells, we further validated the results by perturbing cell-cell contacts in the human breast cancer cell line MCF7A, where depleting E-cadherin by RNAi changed the behaviour from an extensile to a contractile system (Supplementary Figure 5). We then validated that this switch was not a clonal effect by re-expressing E-cadherin which restored collective extensile behaviour to MDCK E-cad KO cells (Supplementary Figure 6a). Moreover, the total defect density within the monolayer of MDCK WT and MDCK E-cad KO cells did not reveal changes in the density of defects between WT and E-cad KO monolayers (Supplementary Figure 6b) indicating that the average distance between the defects and defect-defect interactions are not affected by E-cadherin removal. Furthermore, measuring average flows around −1/2 (trefoil) defects did not show any significant difference between WT and E-cad KO monolayers (Supplementary Figure 2b, and 6c). This is consistent with both simulations (Supplementary Figure 6d) and theories of active nematics,^19,20^ which show that difference in activity affects the self-propulsion of +1/2 defects, while not altering the velocity field of −1/2 defects. Indeed, comparing the Mean-Square-Displacement (MSD) from defect trajectories in WT and E-cad KO monolayers clearly indicates that while +1/2 defects have propulsive behaviour and move faster in WT monolayers, the motion of −1/2 defects is diffusive in both conditions (Supplementary Figure 6e).

We next checked whether the extensile to contractile crossover could be the impact of a change in the behaviour of individual cells. However, based on our traction force data, both single isolated WT and single isolated E-cad KO cells showed contractile behaviour with the forces directed inwards along their elongation axes as cells pulled on the substrate (Figure 1a). This indicates that removing E-cadherin does not change the contractile pattern (intracellular stress) of single cells (Supplementary Figure 7a). Therefore, the change from contractile to extensile behaviour at the collective level can be linked to the presence of E-cadherin which mediates force transmission between neighbouring cells through intercellular interactions.

In order to better discern the competition between intracellular contractile stresses (generated by the actomyosin machinery throughout the cell) and the intercellular stresses (due to neighbour interactions), we varied these two stresses independently using a cell-based model. The model is based on a phase-field formulation^21^ that captures the deformation of individual cells, and has recently been shown to reproduce the formation of topological defects in MDCK monolayers, along with their associated flow field and stress patterns.^22^ In a similar manner as in the experimental analysis, where the orientation of cells were identified through their long axis, in the model a shape tensor, *S*, characterizes the magnitude and direction of cell elongation (Figure 2a). This parameter continuously evolves with the deformation of cells as they push/pull on their neighbours within the monolayer. Following our recent work,^22^ intercellular stresses are defined to be proportional to the shape tensor which allows us to model extensile stresses at the cell-cell contacts (Figure 2a). This form of modeling was inspired by previous studies on adherens junctions and actomyosin interaction which showed that force transduction at the junction can modify the actomyosin network and in turn the cell shape^23^ and was experimentally validated on MDCK WT monolayers through laser ablation experiments where shorter junctions were under higher tension (higher recoil velocity) in comparison to longer junctions which were under lower tension (lower recoil velocity) (Supplementary Figure 3a,b). In addition, an intracellular stress is defined to mimic internal stresses generated by acto-myosin complexes within the individual cells (see Methods for the details of the model). The effect of E-cadherin removal is thus captured in the model by tuning down the intercellular stresses. Just as in the experiments both comet-shaped and trefoil topological defects (+1/2 and −1/2 charges, respectively) are found in the orientation field of the monolayer (Figure 2b) and the average flow fields and strain rate maps around comet shaped defects match those measured for the WT cells (Figure 2c). More importantly, we found that lowering intercellular stresses switched flow direction around comet-shaped topological defects and strain rates in agreement with experimental results of E-cad KO (Figure 2b and c). Quantitative analysis of the simulations showed that reducing the intercellular stresses results in slower dynamics characterized by a smaller root mean square (rms)-velocity (Figure 2d) and generates less correlated patterns of motion characterized by a smaller velocity correlation length (Figure 2e). Moreover, due to the dipolar symmetry of intercellular stresses in the model, simulation results predict that the switch from extensile to contractile behaviour does not alter the isotropic stress patterns, i.e., tension (positive isotropic stress) and compression (negative isotropic stress) around the defects (Figure 3a), when intercellular stresses are reduced.

**Figure 2.**
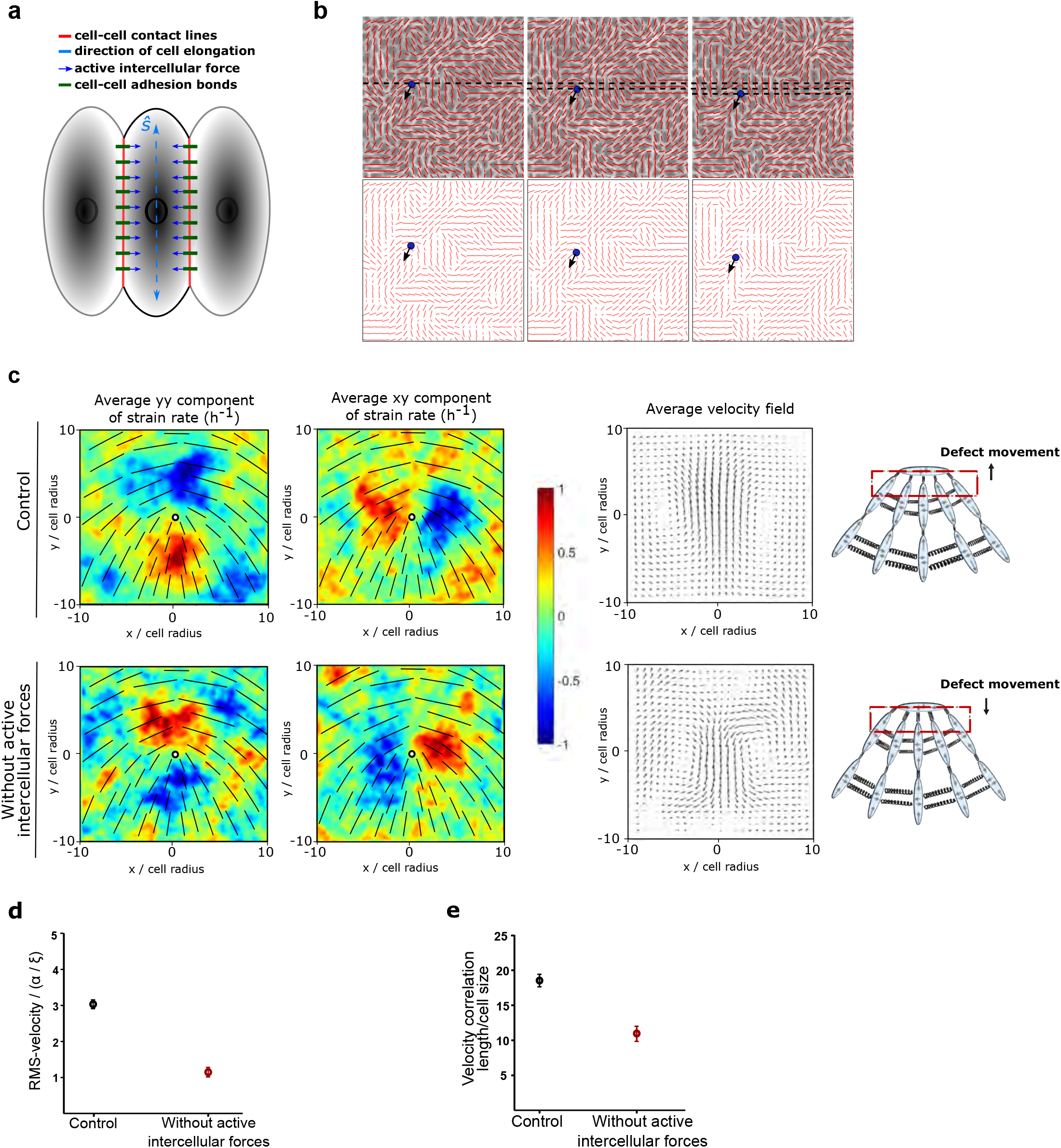
Intercellular stresses change the contractile behaviour of a 2D nematic system. a) Schematic illustrating the model used in numerical simulations which incorporates cell-cell interaction through active intercellular forces. The direction of cell elongation is denoted by the headless vector *ŝ*, which is found from the eigenvector corresponding to the largest eigenvalue of the shape tensor **S** for each cell. b) Numerical simulations for the case without active intercellular stresses, showing: Top, phase contrast images of the cells overlaid with the average local orientation of the cells (red lines). And bottom, average local orientation of the cells (red lines). The blue circle shows the location of a *+1/2* defect and the corresponding arrow indicates the direction of motion of this defect over time. c) Average *yy*- and *xy*-components of strain rate map around *+1/2* defect obtained from simulations (left and middle respectively) and corresponding average velocity flow field (right: *n = 2,083* defects) for the control condition (top) and the condition without active intercellular forces. Colour code is positive for stretching and negative for shrinkage. d) RMS velocity, and e) the velocity correlation length in the monolayer normalized to the individual cell size obtained from *n=30* different simulations for the control condition and the condition without active intercellular forces.

**Figure 3.**
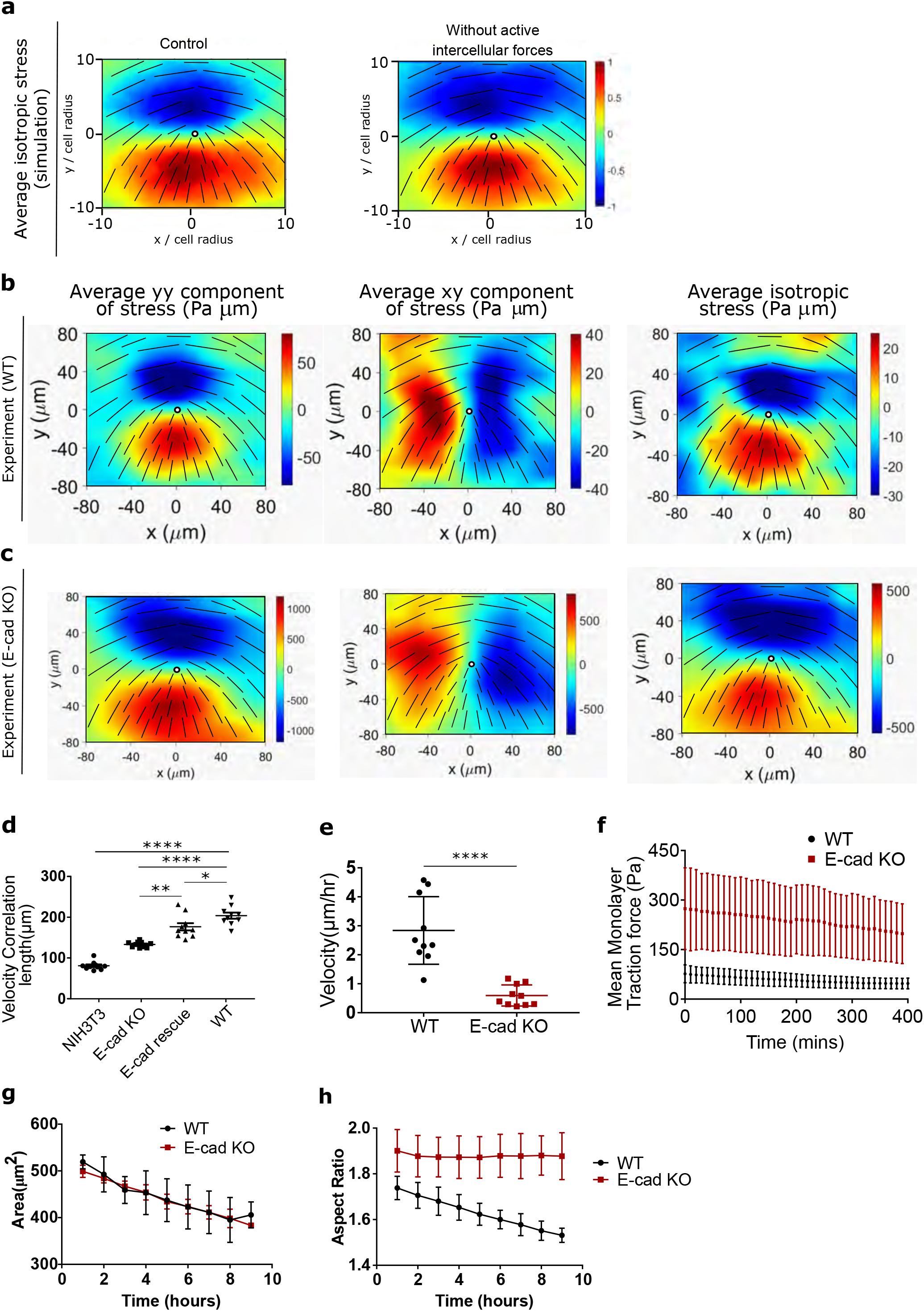
Knocking out E-cadherin increases cell-substrate interactions. a) Average isotropic stress around a *+1/2* defect obtained from simulations for the control condition (left) and condition without intercellular forces (right) (*n = 2,083* defects). b,c) Average *yy* (left)-, *xy* (middle)- and isotropic (right) components of stress around a + 1/2 defect obtained from experiments for (b) MDCK WT (*n = 1,899* defects) and (c) E-cadherin KO (*n = 1,428* defects) from 2 independent experiments. For a and b colour code represents the strength of the stress with positive for tensile state, negative for compression. d, e, f) velocity correlation length (d) (*n=10*), velocity (e) (*n=10*) and mean traction force (f) (*n=12*) of cells within a monolayer for both MDCK WT and MDCK E-cadherin KO cells. g, h) Cell spreading area (g) and aspect ratio (h) of cells within the monolayer obtained from *n=10 different images* for MDCK WT and E-cadherin KO cells as a function of time from *2* independent experiments. The error bars represent the standard deviation. Unpaired t-test was performed resulting in *\*p<0.05, **p<0.01, ***p<0.001* and *\*\*\*\*p<0.0001*.

To test the predictions of the model we first experimentally studied the effect of E-cad KO on the stress patterns around topological defects and collective motion of cells. Using traction force microscopy, we obtain traction forces in the monolayer, from which we infer the associated stress patterns using Bayesian Inversion Stress Microscopy (BISM).^24^ Using a similar approach as strain rate measurements around defects, we are able to compute the average stress fields around comet shaped defects. Our experiments agreed with the simulations in showing no difference in the average isotropic stress patterns around comet shaped defects between the WT (Figure 3b) and E-cad KO monolayers (Figure 3c), while they still show a difference in their flow field (Supplementary Figure 7b),^2^ indicating that the tension and compression around defects are primarily controlled by local cellular organization and elongation, and not the flow field around them. Moreover, measuring the velocity correlation function,^25^ we found it to be consistent with the numerical predictions whereby removing E-cadherin reduces the correlation length compared to the WT monolayers (Figure 3d). This is also in agreement with previous reports which demonstrate a reduction in velocity correlation length of mesenchymal cells with respect to epithelial cells.^25^ Interestingly, by performing rescue experiments to put E-cadherin back, we found an increase in velocity correlation length (Figure 3d) which was very close to that of WT monolayers. This indicates that the perturbation of junctional protein E-cadherin can be used as an effective way of tuning the collective contractility and extensility of the epithelial monolayer.

Comparing the average velocities in the monolayers with and without E-cadherin also agreed with the model’s prediction that the velocity of the monolayer is reduced upon E-cadherin depletion (Figure 3e) at similar density. Interestingly, traction force microscopy measurements revealed that this reduction in velocity is accompanied by a significant (about three fold) increase in the average traction forces that E-cad KO monolayers exert on their underlying substrate in comparison to WT monolayers (Figure 3f). Furthermore, we compared average cell areas within the monolayer for both WT and E-cad KO monolayers and did not notice an appreciable difference in spreading area, although for both WT and E-cad KO monolayers the average cell spreading area reduced over time (Figure 3g). In contrast, the aspect ratio of cells within the WT monolayers reduced over time while the aspect ratio of cells within E-cad KO monolayers did not change over time (Figure 3g). These measurements of velocity reduction, traction force increase, and changes in aspect ratio in the monolayers without E-cadherin, combined together, hinted that the cell-substrate interaction increased as the cell-cell interaction was weakened, indicating a possible cross-talk between intracellular and intercellular interactions as reported previously^26−29^.

To test this, and based on previous studies that showed changes in cellular response to substrate adhesions ^30,31^, we asked if the increase in the average traction force of E-cad KO monolayers was a result of changes in their mechanotransductory response. Using actin staining we first checked for changes in the organization of stress fibers in the cells within a monolayer, as stress fiber formation is an important determinant of force generation by cells on a substrate.^32,33^ Indeed, comparing actin staining of WT and E-cad KO monolayers, we found a considerable increase in stress fibers in the absence of E-cadherin (Figure 4a). Concomitantly, phosphomyosin staining of WT and E-cad KO monolayers showed an increase in the number of phosphomyosin light chain (pMLC2) fibers (Figure 4a) generated at the basal surface within E-cad KO cells. Western blot analyses further revealed an increase in the total level of myosin light chains (MLC2) (Supplementary Figure 4c). Considering these results we reasoned that inhibiting cell contractility in E-cad KO cells may alter their active behaviour. Upon treatment with a mild dose of blebbistatin (5 μM), an inhibitor of Myosin II (Supplementary Figure 8a) E-cad KO monolayers still behave as a contractile system. However, a higher dose (20 μM) of blebbistatin (Supplementary Figure 8b) or 25 μM of Y27632, an inhibitor of ROCK 1 and 2 (Supplementary Figure 8c), resulted in a switch in behaviour from a contractile to that of an extensile system as summarized in Table1. As control, we showed that similar treatments did not affect the extensile behaviour of the WT monolayers (Table 1, Supplementary Figure 8d and e). We then measured the traction forces exerted by cells when treated with 20 μM blebbistatin. As reported previously,^34^ treatment of both WT and E-cad KO monolayers with 20 μM of blebbistatin results in a drastic reduction of traction forces (Figure 4b). This reinforces the importance of cell substrate interaction in dictating the contractile behaviour of E-cad KO monolayers. Thus, removing E-cadherin not only reduces the extensile intercellular stresses, it also increases the intrinsic contractility (intracellular stress) generated by cells at the cell-substrate interface.

**Figure 4.**
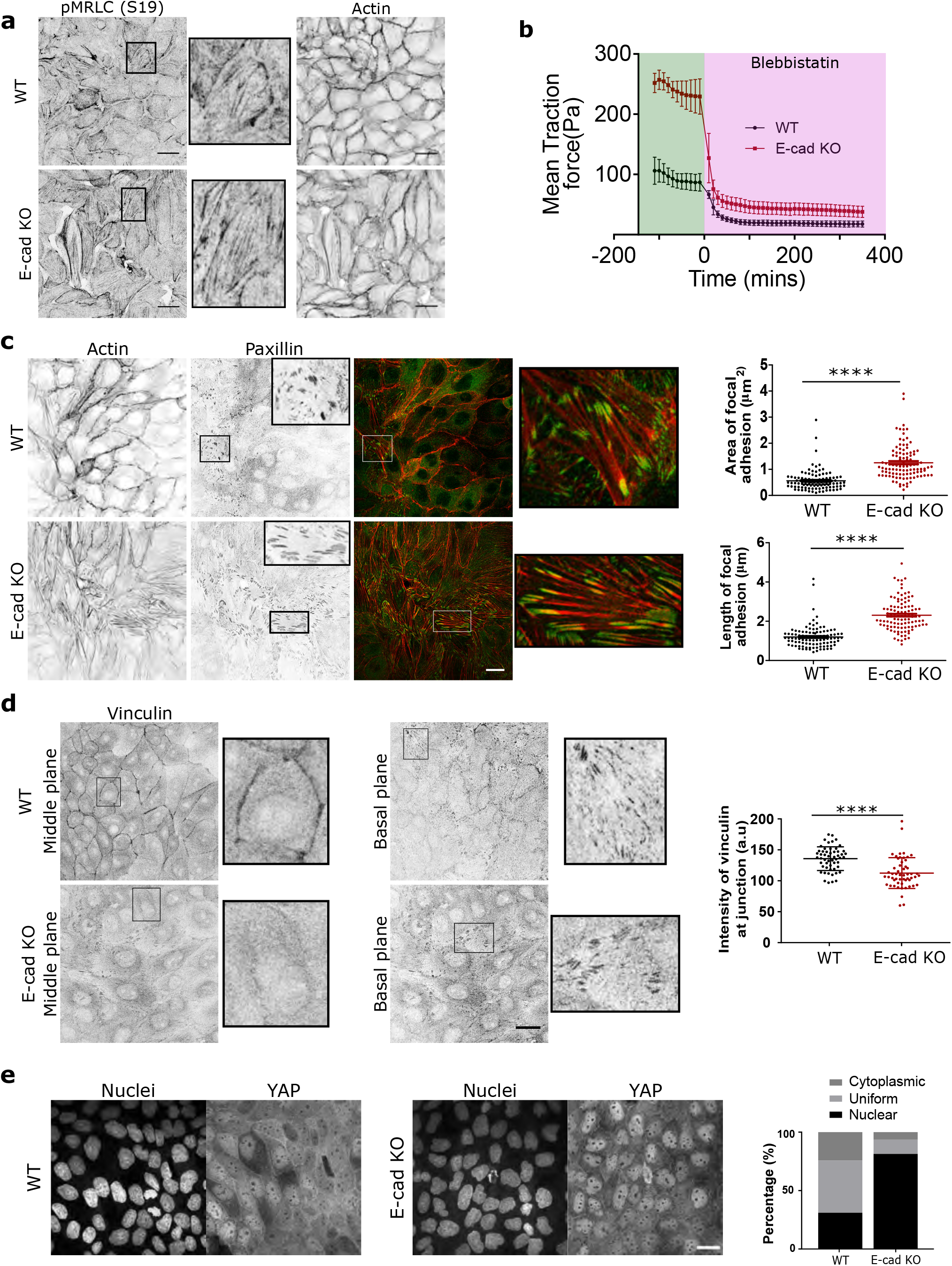
E-cadherin removal triggers mechanotransductive changes within the monolayer. a) pMRLC (left), zoom of pMRLC (middle), actin (right) staining of MDCK WT (top) and E-cadherin KO (bottom) monolayers. b) Evolution of mean traction force of MDCK WT and E-cadherin KO monolayers before and after 20μM blebbistatin treatment (n=10 from 2 independent experiments). c, d, e) actin and paxillin (c), vinculin (d), YAP (green), and nucleus (blue) (e), staining within a monolayer for both MDCK WT and E-cadherin KO cells. c) Area of focal adhesion (left) and length of focal adhesion within the monolayer for *n=106* focal adhesions. d) Mean intensity of vinculin at the cell-cell junction in the middle plane (*n=54*). e) Distribution of YAP in nucleus, cytoplasm, or uniform distribution calculated for *n=1162* cells (MDCK WT) and *n=1008* cells (MDCK E-cadherin KO). Error bars represent the standard deviation. Unpaired t-test was performed leading to *\*p<0.05, **p<0.01, ***p<0.001* and *\*\*\*\*p<0.0001*. Scale bars, *20μm*.

Since focal adhesions (FAs) are known to be mechanosensors at cell-matrix interface,^35^ we then investigated the assembly of FAs in E-cad KO and WT monolayers. By using paxillin staining to determine changes in FAs, we showed a marked increase both in length, and area within the cells (Figure 4c) in the E-cad KO monolayers in comparison to the WT monolayers. More importantly, we found that the E-cad KO modified the subcellular localization of vinculin, a protein which is known to respond and transmit force from both integrin and cadherin based adhesion complexes.^36,37^ While the total level of vinculin remained unchanged in both WT and E-cad KO monolayers (Supplementary Figure 4d), the localization of vinculin was altered, whereby vinculin was mostly present at the cell-cell junctions in WT monolayers, but basally located in E-cad KO monolayers (Figure 4d). We further verified if all paxillin positive focal adhesions were vinculin positive in both WT and E-cad KO monolayers and observed a strong correlation between them (Pearson’s coefficient of 0.8842 and 0.8843 for WT and E-cad KO) as shown in Supplementary Figure 9, reiterating our observed increase in cell-substrate interaction in the absence of E-cadherin.

Since Yes-associated protein (YAP) transcriptional activity is also known to modify cell mechanics, force development and FA strength,^38,39^ we investigated the localization of YAP within E-cad KO monolayers. Interestingly, we found that YAP was predominantly localized to the nucleus in E-cadherin KO monolayers (Figure 4e), which corresponds to the active state of YAP. This is in agreement with previous studies that reported an activation of YAP through nuclear accumulation in the absence of E-cadherin or in well spread cells.^30,40,41^ Taken together, our results show that removing E-cadherin enhances the formation of stress fibers, promotes YAP activation, alters viculin localization, and leads to a marked increase in the formation of focal adhesions and their linkage to the substrate, in turn triggering a contractile behaviour.

Our force measurements together with acto-myosin activity and adhesion patterns establish that the extensile or contractile nature of epithelial cells at a collective level relies on the interplay between active stresses at cell-cell and cell-matrix interfaces. To further explore this crossover we plated cells on a soft (2.3 kPa) polyacrylamide (PA) gels, recalling that cellular responses on soft substrates leads to lower contractility and less stable focal adhesions.^42^ MDCK WT monolayers remained extensile regardless of substrate stiffness (Supplementary Figure 10a), while E-cad KO cells switched from contractile to extensile behaviour on a soft substrate (around 2.3 kPa) (Supplementary Figure 10b). Taken together, these experiments show that tuning cell-cell and cell-substrate adhesion can result in a switch between extensile and contractile behaviour of cell monolayers further validating our observatin that blebbistatin treatment drastically reduced traction forces (Figure 4d) and switched the behaviour of E-cad KO monolayers from contractile to extensile. It is possible in the simulations to further explore this crossover by continuously varying the strength of intra- and inter-cellular stresses, independently. The results are summarized in a stablity-phase diagram that classifies the monolayer behaviour as extensile or contractile based on the direction of the defect motion (Supplementary Figure 11a). The non-symmetric structure of the stability-diagram further highlights the different impacts of intra- and inter-cellular stresses on the direction of defect motion. In our simluations, while intracellular stresses act within single cells and are along the direction of cell polarity, the intercellular stresses arise in between neighboring cells and are proportional to the cell deformation. As such, intercellular stresses can reinforce themselves: small cell deformations lead to intercellular stresses that further enhance cell deformation, generating stronger intercellular stresses. We conjecture that this bootstrap mechanism results in intercellular stresses to more strongly affect the collective behavior of the monolayer compared to their intracellular counterparts.

Based on this difference in contractile and extensile behaviour we then used the model to simulate the interaction between the extensile and contractile systems. The results showed that cells were able to separate out into two different phases over time when mixed at 50-50 ratio (Figure 5a and Supplementary Figure 11b, Video 3), where extensile cells were surrounded by contractile ones. We were able to replicate this experimentally (Figure 5b and Supplementary Figure 11c, Video 4) whereby WT and E-cad KO cells separate out into two different phases with WT (extensile) cells surrounded by E-cad KO (contractile) cells when plated at a 50-50 ratio (Figure 5a). While thermodynamic mechanisms such as differential adhesion and difference in line tension between two cell types have been shown to govern phase separation in 3D cell aggregates,^43–45^ active cell sorting in monolayers with strong substrate adhesion, has not been directly observed to the best of our knowledge. We, therefore, sought to further explore the possible distinctions between the cell sorting, as observed here, and the well-established differential adhesion and differential line tension hypotheses. To this end, we first quantified the degree of phase separation by measuring the mixing-index of a mixture of WT and E-cad KO cells defined as the number of homotypic neighbours over the total number of cells.^46,47^ In the segregation mechanism based on differential line tension this mixing-index grows with a power-law exponent with time and approaches one.^47^ However, as evident from both experiments and simulations, the mixing-index in our system saturates and complete phase separation is never obtained (Figure 5a and b). We conjecture that this is partly because of strong cell-substrate adhesion that dominates over any possible difference in line tensions and also due to a fundamental difference between activity-driven phase separation and thermodynamic mechanisms. In addition, phase separation based on differential line tension posits that – independent of the asymmetry of the binary mixture - the phase with higher line tension always forms aggregates that are enveloped by the cells with lower line tension to minimize the free energy of the mixture.^43,45^

**Figure 5.**
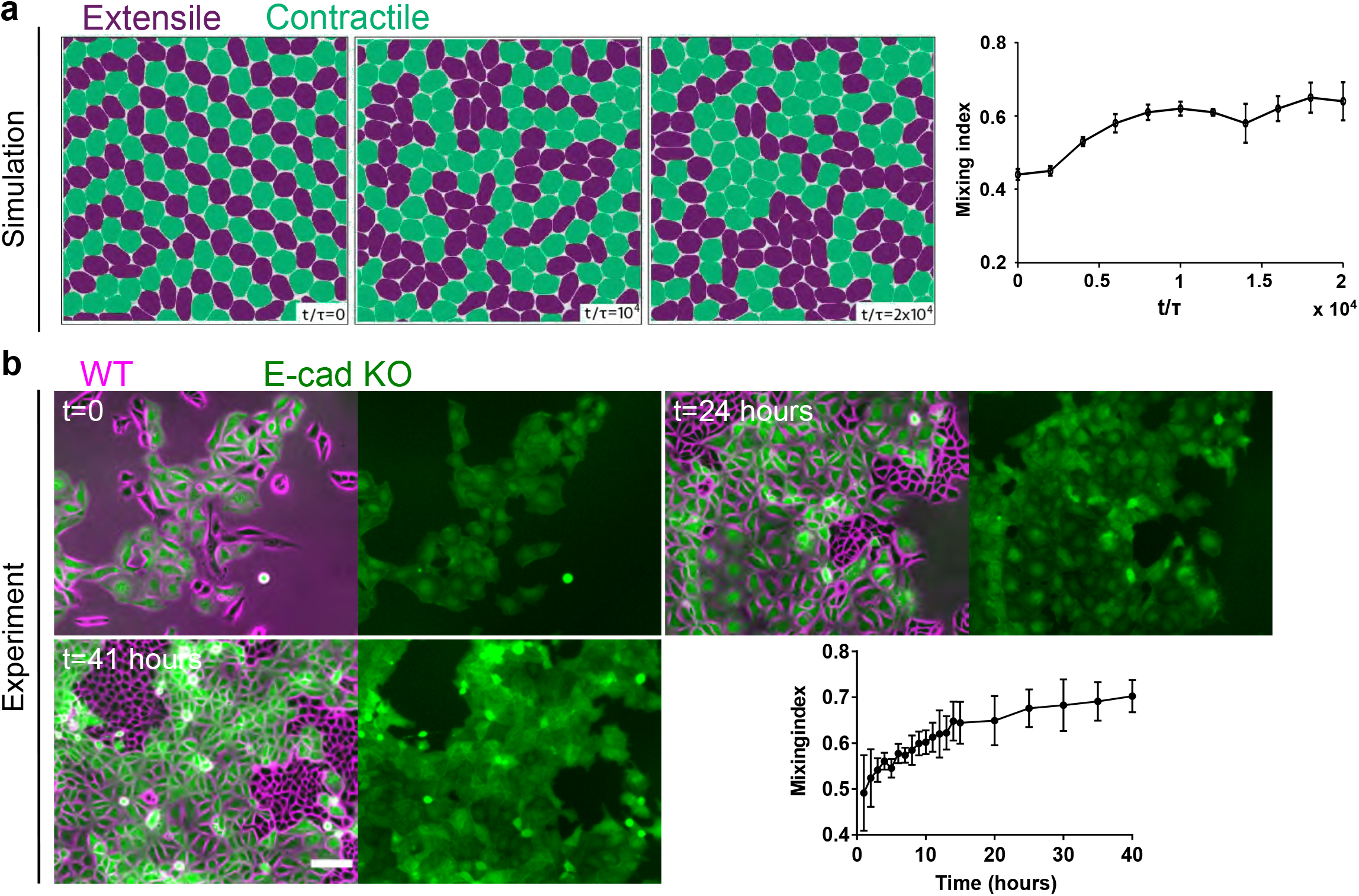
Cell sorting triggered by change in nematic behaviour of monolayers. a,b) Time lapse sorting of extensile and contractile cells observed over time represented by mixing index in simulations (a) and experiments (b) of MDCK WT (magenta) and E-cadherin KO cells tagged with LifeAct GFP (green). In (a) ζ_s_/Rα = 0.042, ζ_Q_/Rα = −0.062 for the extensile cells and ζ_s_/Rα = 0.0, ζ_Q_/Rα = −0.062 for the contractile cells. Mixing index was obtained from two independent simulations and the error bars mark the standard deviation. Mixing index in experiments (b) was obtained from n=5 different clusters from 2 independent samples. c, d, e, f, g) Demixing (left) and isotropic stress field (right) for different conditions tested in the simulations, line tension (c), steric repulsion (d), adhesion (e), activity based extensile and contractile (f) and experimentally obtained phase separation (left) and isotropic stress field (right). Error bars represent the standard deviation. Scale bars: *100μm*.

To test this, we performed mixing experiments by varying the percentage of WT versus E-cad KO cells, (30/70 and 70/30, respectively; Figure 6a and Supplementary Figure 11d,e). In the latter case, we could even observe E-cad KO colonies surrounded by WT cells which could not be simply explained by the differential adhesion hypothesis and was not observed in previous adhesion based studies governed by cortical/line tension.^43,45−49^ We were able to replicate this in our simulations (Figure 6b). Moreover, to further test the unmixing phase we thought to probe the unmixing of two cell types with and without E-cadherin, but both showing extensile behaviour. Since 20 μM blebbistatin was shown to reverse the contractile behaviour of E-cad KO monolayers from contractile to extensile (Table 1 and Supplementary Figure 8), we treated a mixture of WT and E-cad KO plated at 50/50 ratio with blebbistatin after unmixing. Upon blebbistatin treatment, we see a drop in the mixing index (Figure 6c, Video 5). In addition, the clear boundaries formed in an untreated sample were lost characterized by the loss of circularity of WT colonies upon blebbistatin treatment (Figure 6c).

**Figure 6.**
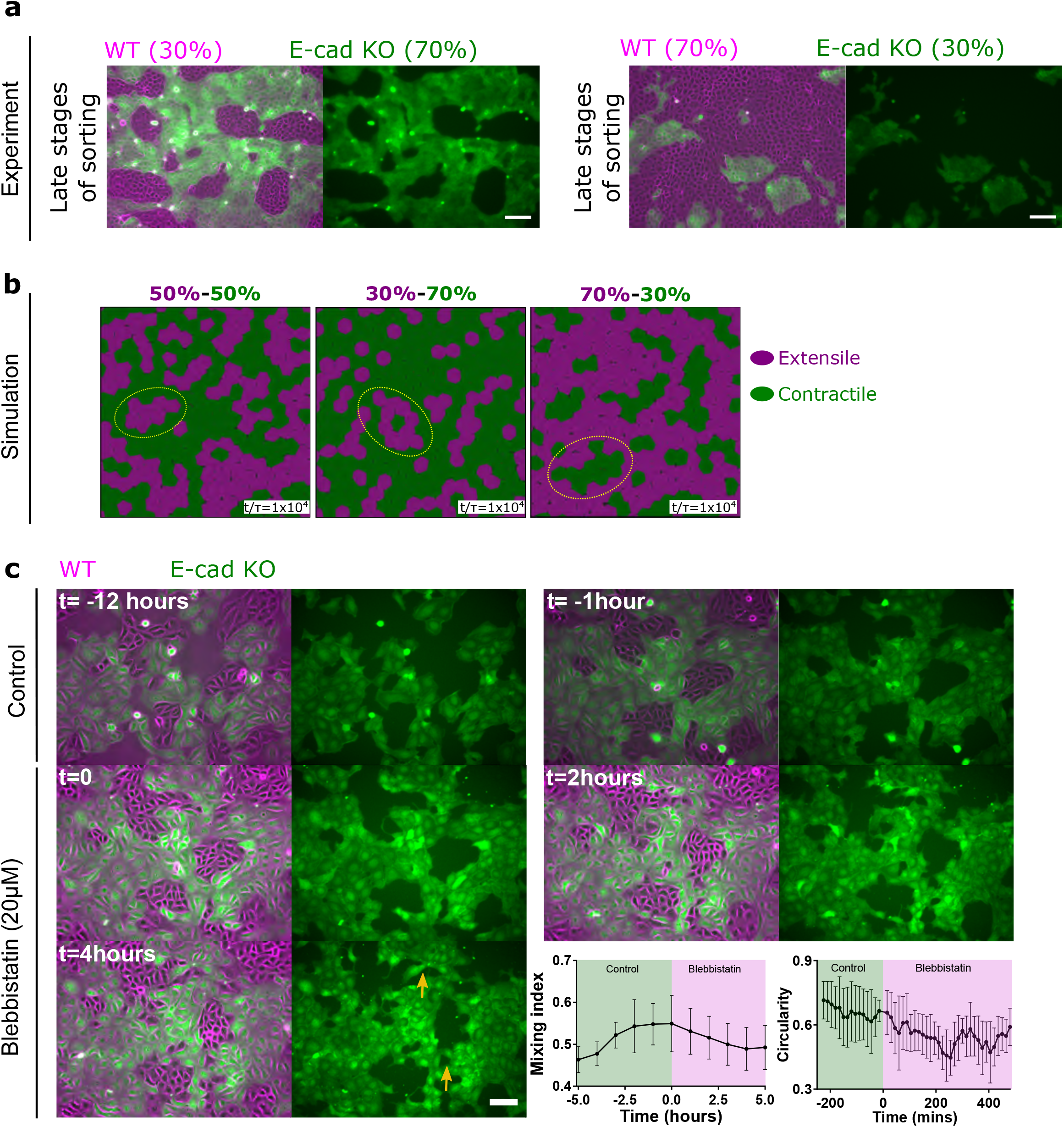
Cell sorting is governed by activity of the system. a) Demixing of MDCK WT and E-cadherin KO at different starting densities, WT (30%) and E-cadherin KO (70%) (left) and WT (70%) and E-cadherin KO (30%) (right). b) Demixing of extensile and contractile particles obtained from simulations at different starting densities. Extensile and contractile particles are mixed at 50-50 (left), 30-70 (middle) and 70-30 (right) respectively. In (b) ζ_s_/Rα = 0.016, ζ_Q_/Rα = −0.016 for the extensile cells and ζ_s_/Rα = 0.0, ζ_Q_/Rα = −0.016 for the contractile cells. c) Demixing phase observed before and after the addition of 20μM blebbistatin characterized by mixing index (left) (n=5) and circularity of several colonies (right) (n=5). Error bars represent the standard deviation. Scale bars: *100μm*.

Taken together, these results reinforce the fundamental distinctions between phase separation in systems with differences in activity in comparison to well-established differential line tension or differential adhesion mechanisms. Even though tissue segregation was first exemplified based on differences in cadherin-mediated surface tension,^43,48,50^ it was later shown that intercellular adhesion is not the only mechanism that triggers cell sorting.^51^ Theoretical predictions have suggested that cell sorting could be driven by a combination of cell surface tension and contractility.^45,46^ While, we cannot completely rule out the contribution of differential adhesion or differential line tension towards the sorting between WT and E-cad KO cells, our results clearly demonstrate the importance of cell-substrate interaction and intracellular stresses as key regulators of cell sorting in cellular monolayers with strong adhesion to substrate.

The results presented in this work show that epithelial cells are able to maintain their collective behaviour through a coordination of intercellular and intracellular stresses. Intercellular stresses are mediated through adherens junctions, while intracellular stresses could be mediated through changes in substrate interaction and actomyosin machinery. Using a combination of in-silico modelling and extensive experimental studies we have shown that perturbation of E-cadherin in MDCK cells, increases their substrate interaction in addition to changing their active nematic behaviour from extensile (WT) to contractile (E-cad KO) similar to a monolayer of fibroblast which behaves as a contractile unit. Our experimental results also show that perturbation of adherens junctions are accompanied by molecular level changes, including reduced levels of vinculin at cell-cell contacts, together with an increase in focal adhesion size and area in the absence of E-cadherin, and increase in the number of actin stress fibers on the basal layer. While, using our numerical model we were able to study how varying inter and intracellular stresses impacts the active behaviour of cells. In addition, mixing the two different systems revealed that these differences in active behaviour were sufficient to drive sorting of these domains into an unmixed phase over time. Comparing our observations of sorting with previously observed studies and hypothesis^43–45^ highlights fundamental distinctions that arise due to the difference in the nature of active forces. These observations bring in a new understanding to the existing models of differential adhesion. Having understood the role of extensility and contractility in dictating demixing (sorting) of cells, this approach could be expanded to studying other biological processes such as tissue growth, development and tissue homeostasis. For instance, recent studies demonstrated the importance of nematic organization of actin cytoskeleton in Hydra during morphogenesis,^52^ while other studies have begun to explore the role of liquid-crystal ordering during morphogenesis^53^ and *in – vivo* epithelial tissue patterning.^54^ These findings highlight the importance of active nematic behaviours at a collective level to understand tissue shape and organization, factors central to morphogenesis.^52,53,55−58^ As such, the adaptation of cellular systems from extensile to contractile behaviours might be a crucial mechanism by which a collective living system undergoes morphological changes (sorting or tissue organization) based on a transition from a cohesive to a less coordinated organization. Such a transition relying on the cross-talk between cell-cell and cell-matrix interactions may provide a new mechanism to understand cell migration during development, wound healing, and collective cancer cell invasion.

## Acknowledgements

This work was supported by the European Research Council (Grant No. CoG-617233), LABEX Who Am I? (ANR-11-LABX-0071), the Ligue Contre le Cancer (Equipe labellisée), and the Agence Nationale de la Recherche (‘POLCAM’ (ANR-17-CE13-0013 and ‘MechanoAdipo’ ANR-17-CE13-0012). We acknowledge the ImagoSeine core facility of the IJM, member of IBiSA and France-BioImaging (ANR-10-INBS-04) infrastructures. A. D. acknowledges support from the Novo Nordisk Foundation (grant No. NNF18SA0035142), Villum Fonden (Grant no. 29476), and funding from the European Union’s Horizon 2020 research and innovation program under the MarieSklodowska-Curie grant agreement No. 847523 (INTERACTIONS). LB has received funding from the European Union’s Horizon 2020 research and innovation programme (Marie Sklodowska-Curie grant agreement 665850-INSPIRE). T.B.S. acknowledges support from the Lee Kuan Yew (LKY) Postdoctoral fellowship and Singapore Ministry of Education Academic Research Fund Tier 1 (R-397-000-320-114). S.G and A. Y were supported by project grants and fellowships from the National Health and Medical Research Council of Australia (1123816 and 1139592) and Australian Research Council (DP190102871). We would like to thank Phillipe Marcq for help with implementation of BISM code, Marina A. Glukhova for providing the vinculin antibody and Sylvie Robine for the ZO1 antibody. AD and JMY acknowledge Guanming Zhang for helpful discussions regarding the model. We also thank the members of cell adhesion and mechanics team at Institut Jacques Monod, Matthieu Piel and Francois Gallet for insightful discussions.

## Author contributions

L.B, T.B.S, A.D, J.Y, R.M.M and B.L designed the research. G.H.N.S.N and T.D developed the MDCK E-cadherin KO cell line. L.B performed experiments and analysed the results. S.S helped in the PA gel experiments. M.T performed and quantified laser ablation experiments. S.G performed the MCF7A experiments. A.D, R.M, performed the in silico simulations. T.B.S contributed to the analysis tools. A.S.Y, Y.T, R.M.M, A.D, J.Y, and B.L supervized the project.

## Code Availability

Nematic analysis was performed using a custom made code which has been published in Saw et al, Nature 2017. This is available upon request. Numerical analyses were performed using a custom made code “CELADRO: Cells as Active Droplets”, which is an open source code that we have deposited on GitHub (https://github.com/rhomu/celadro). All the analysis tools are available upon request.

