## Supplementary Figure 1 for "Nature of active forces in tissues: how contractile cells can form extensile monolayers"

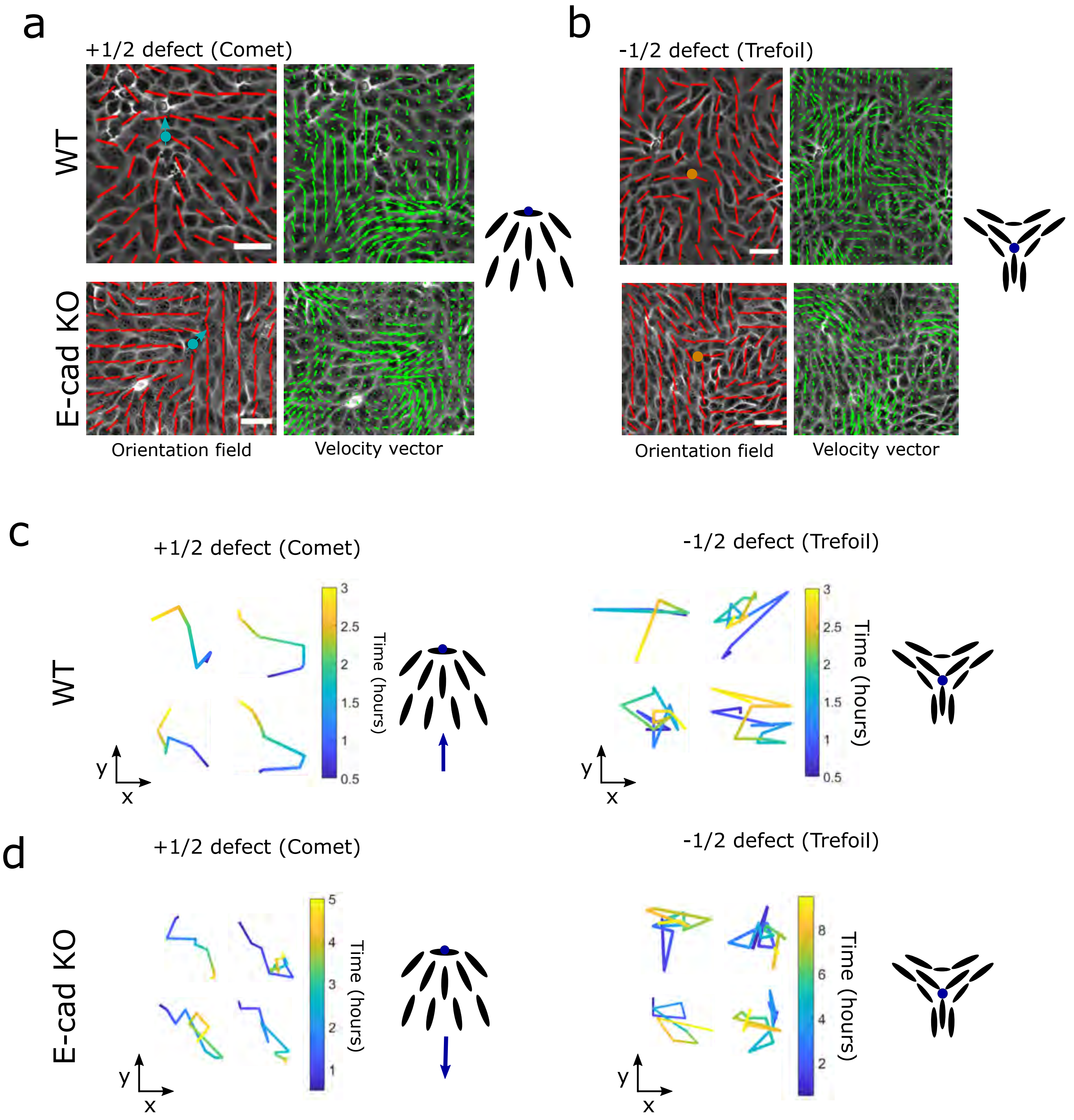


**Figure S1| MDCK WT behave as an extensile system and MDCK E-cadherin KO behave as a contractile system.** a, b) Orientation field (left) and velocity vectors (right) around a single comet shaped (+1/2) defect (a) and trefoil (-1/2) defect obtained from WT (top) and E-cadherin KO (bottom) monolayers. c, d) Trajectory of several comet (+1/2) (left) and trefoil (-1/2) (right) shaped defects obtained from MDCK WT (c) and MDCK E-cadherin KO (d) monolayers. Scale bars: 40µm.
