## Supplementary Figure 2 for "Nature of active forces in tissues: how contractile cells can form extensile monolayers"

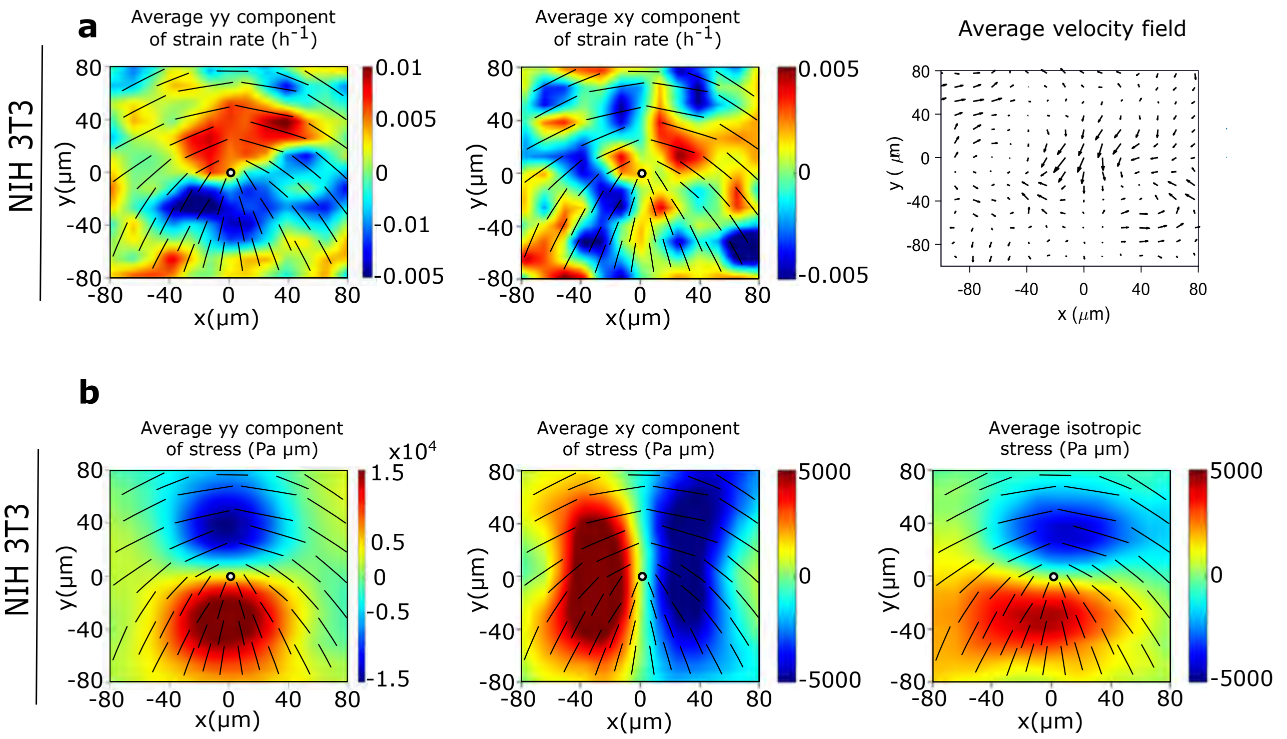


**Figure S2| Fibroblasts behave as a 2D contractile active nematic.** a) Average yy- and xy-components of strain rate map around +1/2 defect obtained from experiments (left and middle respectively) and corresponding average flow field (right) (n = 1489 defects from 2 independent experiments) for NIH3T3 cells. Colour code is positive for stretching and negative for shrinkage. b) Average yy (left)-, xy (middle)- and isotropic (right) components of stress around a +1/2 defect obtained from experiments for NIH3T3 (n = 1,428 defects from 2 independent experiments).
