## Supplementary Figure 3 for "Nature of active forces in tissues: how contractile cells can form extensile monolayers"

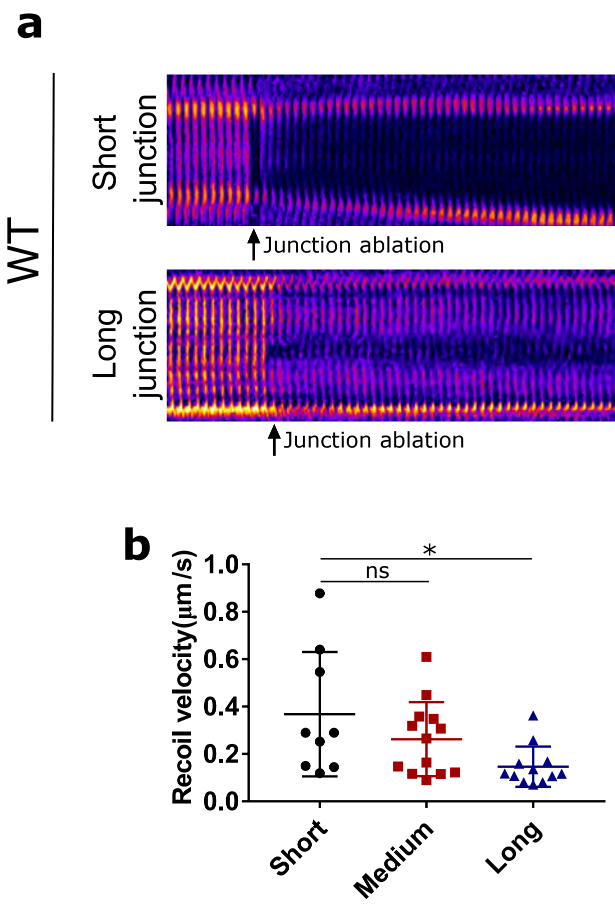


**Figure S3| MDCK WT behave as an extensile system.** a) Kymograph of a short junction (<10μm) (top) and long junction (>15μm) (bottom) before and after laser ablation.  B) Recoil velocity after laser ablation for short (<10μm) (n=9) (N=4), medium (10-15μm) (n=13)(N=4) and long junctions (> 15μm) (n=12) (N=6). n, is the number of junctions ablated and N is the number of independent experiments from which these results were obtained. Error bars represent the standard deviation. ANOVA test was performed leading to *p<0.05, **p<0.01, ***p<0.001 and ****p<0.0001. Scale bars, 20μm.
