## Supplementary Figure 4 for "Nature of active forces in tissues: how contractile cells can form extensile monolayers"

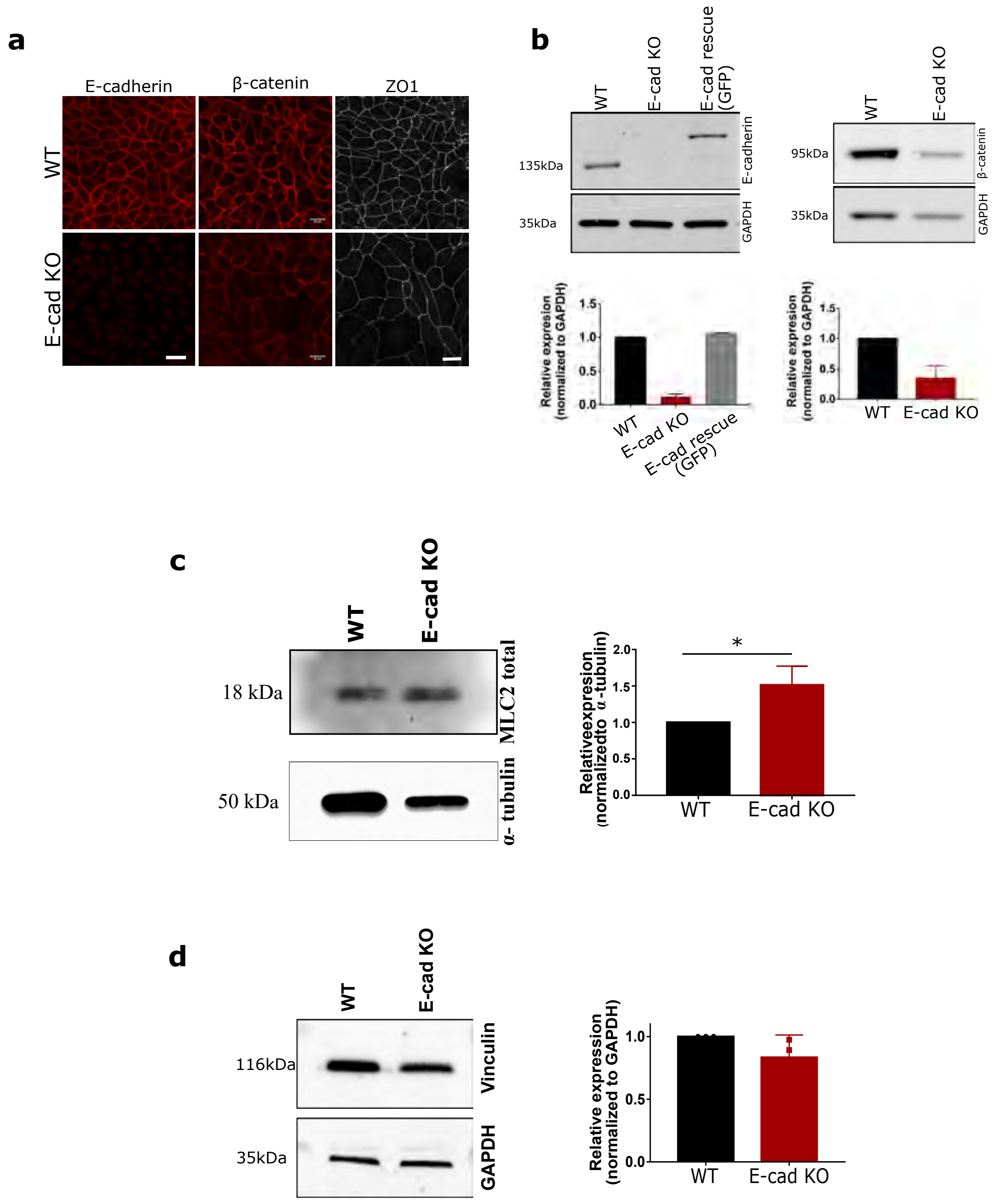


**Figure S4| Characterization of MDCK E-cadherin KO cells.** a) Immunofluorescence staining (top) of E-cadherin (left), β-catenin (middle) and ZO1 (right), along with a b) representative western blot and quantification for E-cadherin (left) (n=3) and β-catenin (right) (n=3). Scale bars, 20μm. c) Western blot analysis of total MLC and quantification from 3 independent experiments normalized to α-tubulin. d) Western blot analysis of vinculin and quantification from 3 independent experiments normalized to GAPDH. Error bars represent the standard deviation.
