## Supplementary Figure 5 for "Nature of active forces in tissues: how contractile cells can form extensile monolayers"

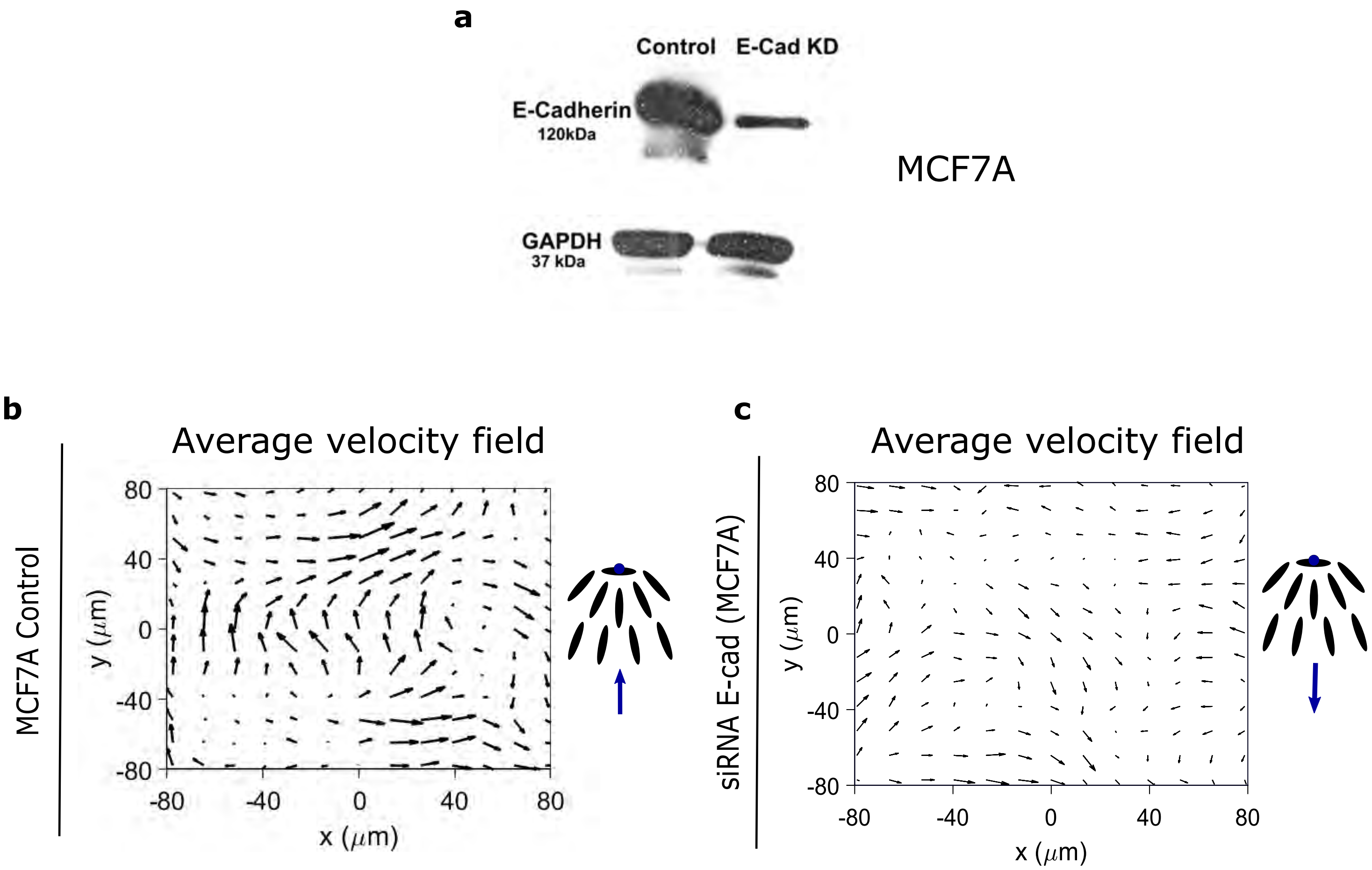


**Figure S5| Contractile and extensile behaviour of MCF7A cells.** a) Western blot showing the reduced level of E-cadherin in siRNA generated E-cadherin KD cell line for MCF7A cells. b and c) Average flow field for MCF7A control cells (n = 2047 defects from 3 independent experiments) (b) and siRNA E-cadherin KD MCF7A cells (n = 1256 defects from 3 independent experiments) (c).
