## Supplementary Figure 6 for "Nature of active forces in tissues: how contractile cells can form extensile monolayers"

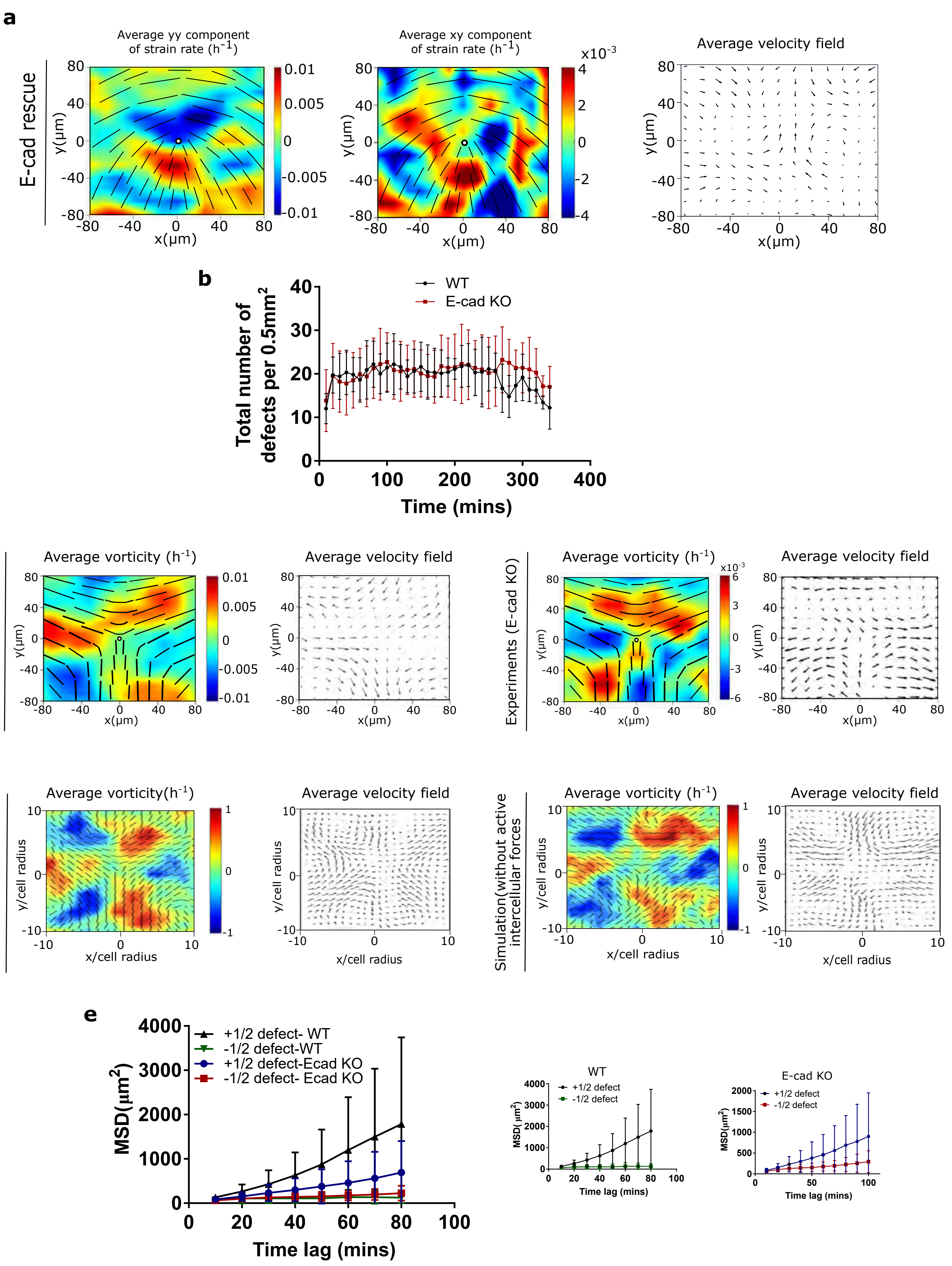

**Figure S6| E-cadherin rescue changes the behaviour to a 2D extensile active nematic liquid crystal and flow field around trefoil shaped defects.** a) Average *yy*- and *xy*-components of strain rate map around comet (+*1/2)* defect obtained from experiments (left and middle respectively) and corresponding average flow field (right) (*n* = *1767* defects from *2* independent experiments) for MDCK E-cadherin KO cells rescued with E-cadherin GFP. b) Total number of defects obtained per $0.55mm^2$ as a function of time on MDCK WT and MDCK E-cadherin KO monolayers. (n=10) from 2 independent experiments. c) Average vorticity and velocity field around trefoil (-1/2) defects in WT (left) (n=1934) and E-cadherin KO (right) (n=2028) monolayers. d) Average vorticity and velocity field around trefoil (-1/2) defects in control (left) (n=3200) and condition without active intercellular forces (right)(n=3200) monolayers obtained from simulations. e) Mean square displacement (MSD) plotted against time lag for comet (+1/2) and trefoil (-1/2) defects obtained from MDCK WT and MDCK E-cadherin KO monolayers (n=11). Error bars represent the standard deviation.
