## Supplementary Figure 7 for "Nature of active forces in tissues: how contractile cells can form extensile monolayers"

a

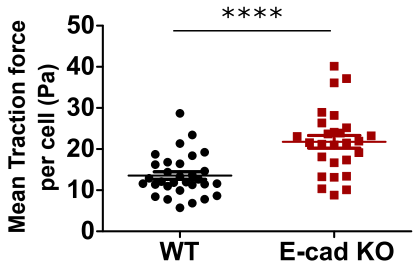


b

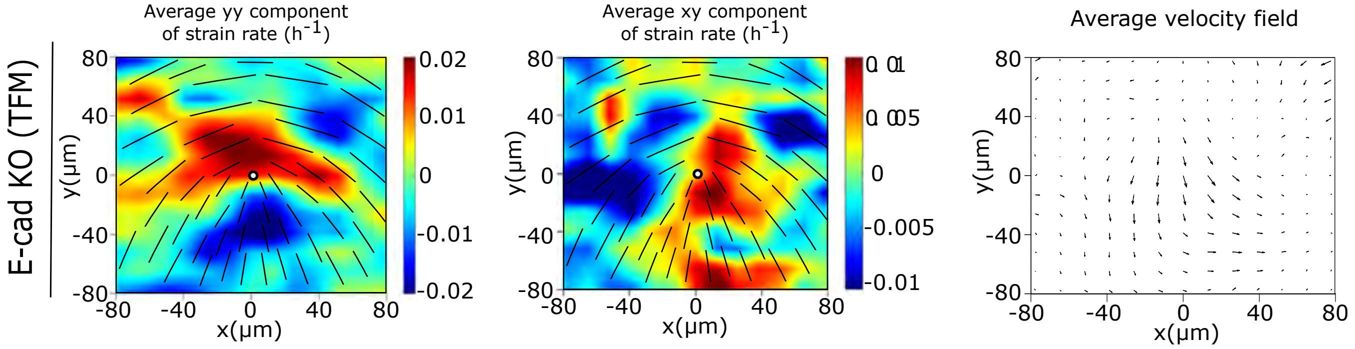


**Figure S7| E-cadherin removal does not affect the contractile behaviour of single cells** a) Mean traction force for both MDCK WT (n=31) and MDCK E-cadherin KO cells (n=27). b) Average yy- and xy-components of strain rate map around comet (+1/2) defect obtained from experiments (left and middle respectively) and corresponding average velocity flow field (right) (n = 1428 defects from 2 independent experiments) for MDCK E-cadherin KO cells plated on PDMS substrates of stiffness 15kPa from which stress maps were obtained in Figure 3b’.
