## Supplementary Figure 8 for "Nature of active forces in tissues: how contractile cells can form extensile monolayers"

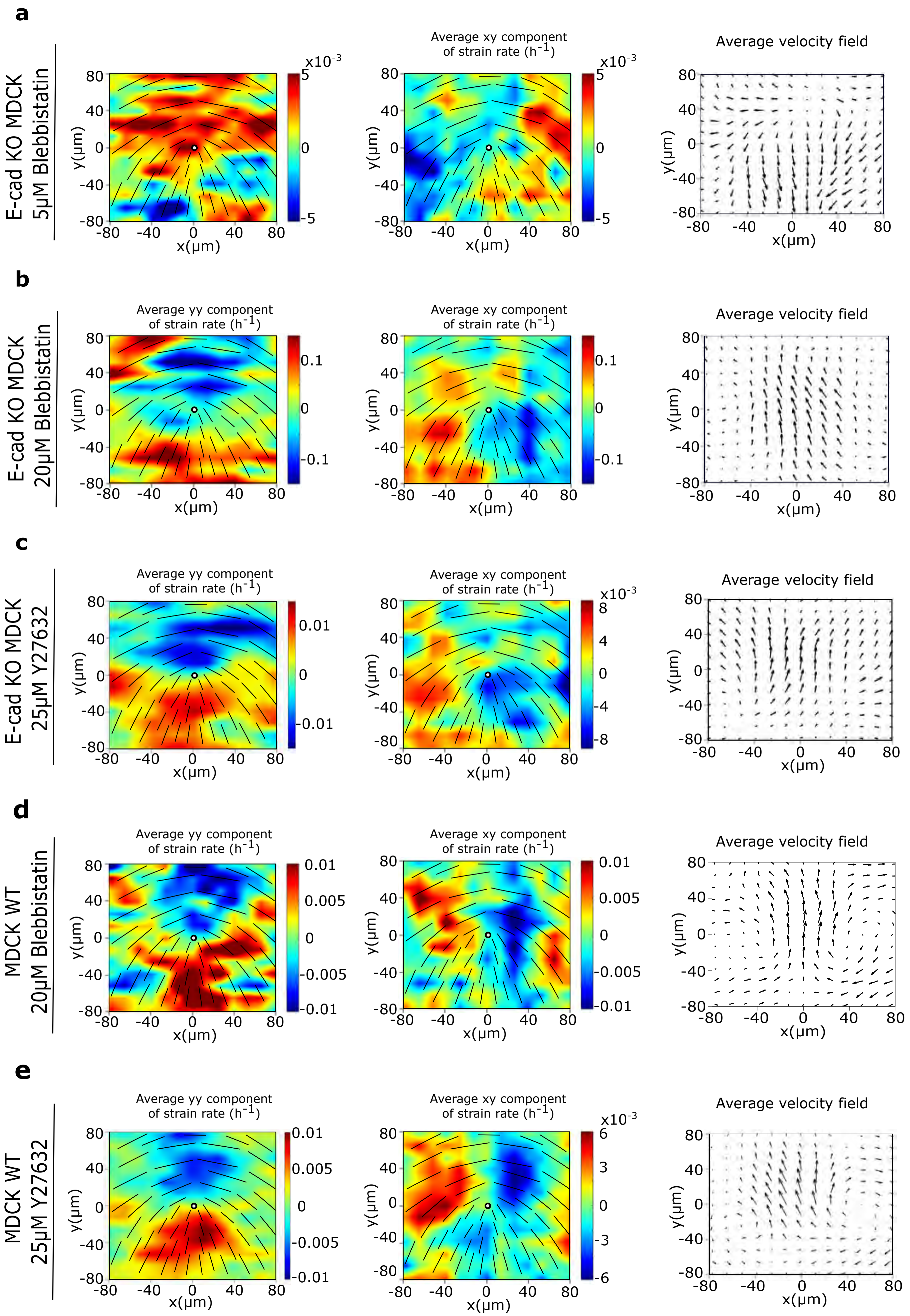

**Figure S8| Drug treatment changes 2D active nematic behaviour of MDCK E-cadherin KO cells.** a, b, c) Average *yy*- and *xy*-components of strain rate map around *+1/2* defect obtained from experiments (left and middle respectively) and corresponding average velocity flow field (right) for MDCK E-cadherin KO cells treated with *5μM* blebbistatin (a) (*n = 2174* defects from 2 independent experiments), *20μM* blebbistatin (b) (*n = 1223* defects from 2 independent experiments), and *25μM* Y27632 (c) (n = 1965 defects from 2 independent experiments). d, e) Average *yy*- and *xy*-components of strain rate map around *+1/2* defect obtained from experiments (left and middle respectively) and corresponding average velocity flow field (right) for MDCK WT cells treated with *20μM* blebbistatin (d) (*n = 1287* defects from *2* independent experiments), and *25μM* Y27632 (e) (*n = 2472* defects from *2* independent experiments).
