## Supplementary Figure 9 for "Nature of active forces in tissues: how contractile cells can form extensile monolayers"

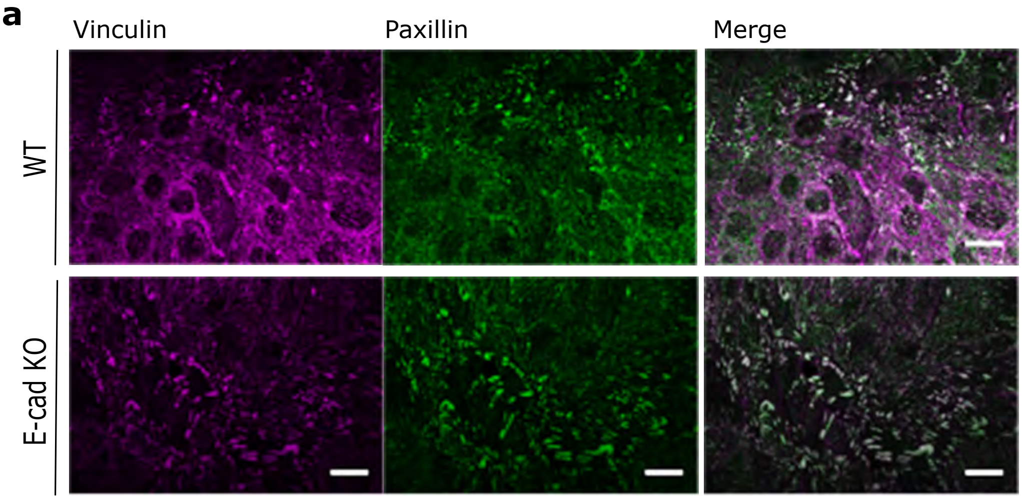


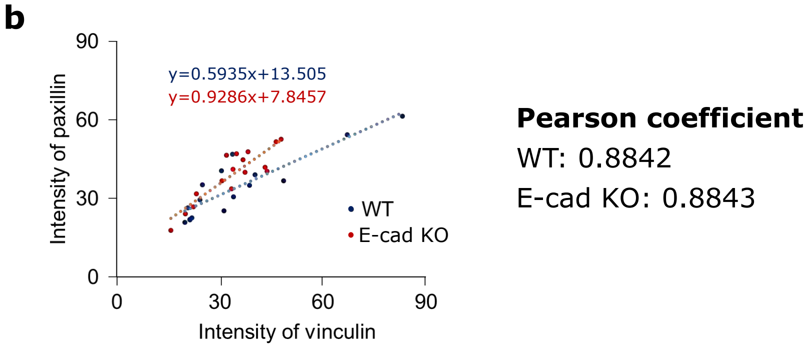


**Figure S9| Colocalization of vinculin and paxillin.** a) Immunostaining of basal plane of vinculin (left), paxillin (middle) and merge (right) in MDCK WT (top) and MDCK E-cadherin KO monolayers. b) Intensity of vinculin plotted against paxillin for n=15 focal adhesions in MDCK WT and n=16 focal adhesions in MDCK E-cadherin KO monolayers. Scale bars: 20μm.
