## Supplementary Figure 10 for "Nature of active forces in tissues: how contractile cells can form extensile monolayers"

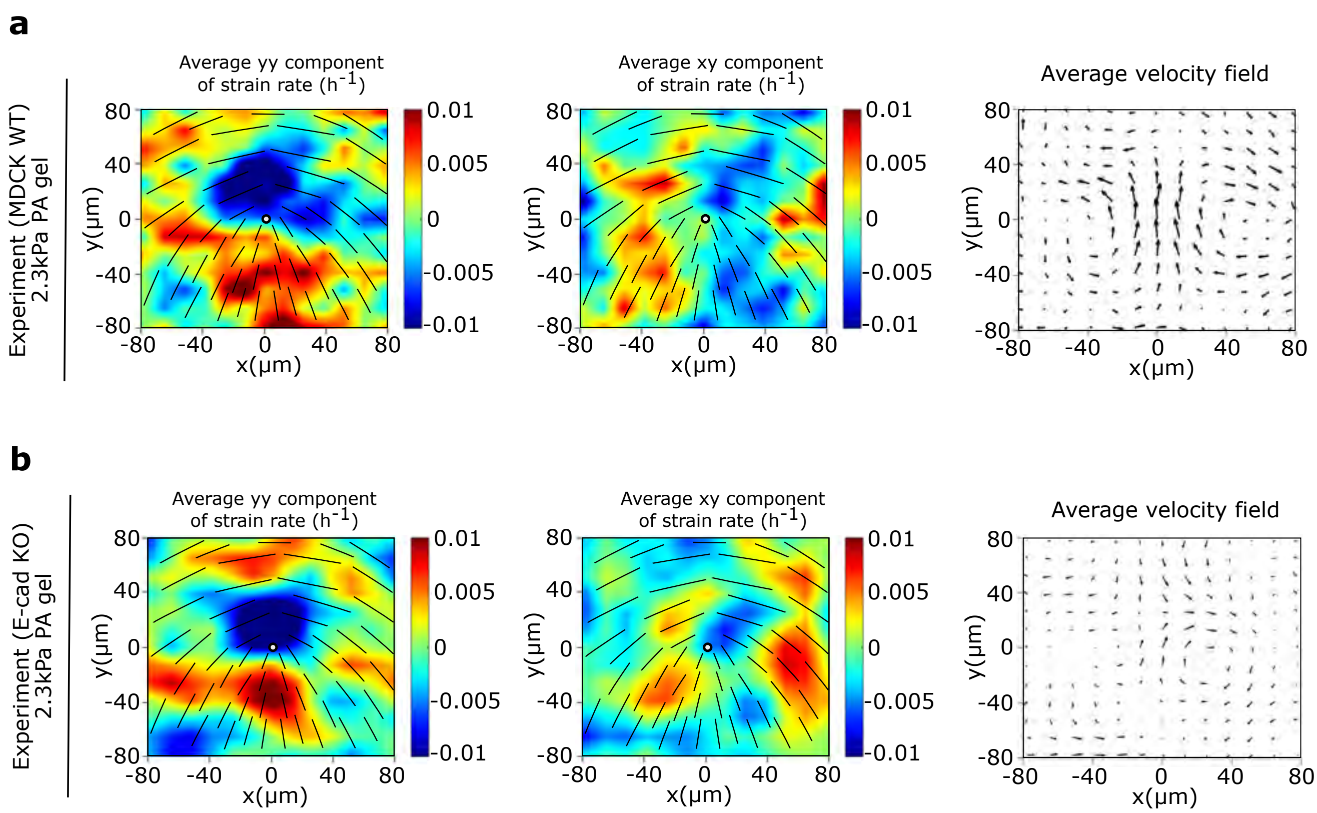


**Figure S10| Substrate rigidity alters E-cadherin KO behaviour.** a, b) Average yy- and xy-components of strain rate map around +1/2 defect obtained from experiments (left and middle respectively) and corresponding average flow field (right) for MDCK WT cells (a) (n = 1426 defects from 2 independent experiments) and E-cadherin KO cells (b) (n = 1041 defects from 2 independent experiments).
