## Supplementary Figure 11 for "Nature of active forces in tissues: how contractile cells can form extensile monolayers"

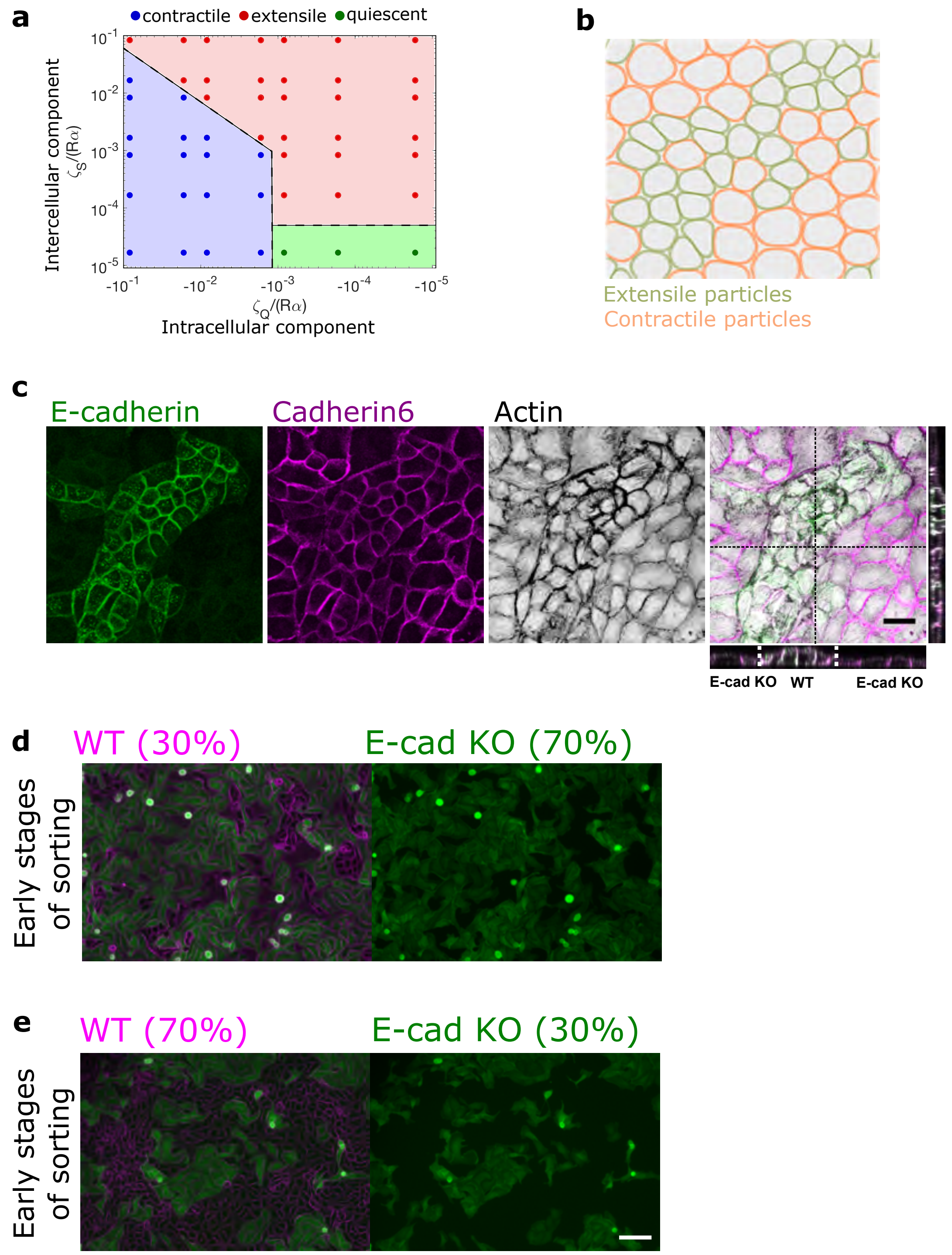

**Figure S11| Phase diagram on activity change and activity based cell sorting in a mixed culture of MDCK WT and MDCK E-cadherin KO.** a) Phase diagram showing the transition of extensile and contractile behaviour with varying values of intercellular and intracellular stresses obtained from simulations. b) Phase separation (demixing) observed from simulations where the contractile particles (orange) are surrounded by extensile particles (green). c) Cell sorting (demixing) observed for a mixture of MDCK WT and MDCK E-cadherin KO cells, where WT cells are surrounded by E-cadherin KO cells (E-cadherin, green, cadherin 6, red, actin, black). XZ and YZ projection show the height difference between the two cells when mixed. Scale bars, *20μm*. d,e) Early stages of cell sorting when MDCK WT (magenta) and MDCK E-cadherin KO (green) monolayers are mixed at 30-70 (d) and 70-30 (e) ratio. Scale bars: *100μm*.
