## Supplementary material for "Nature of active forces in tissues: how contractile cells can form extensile monolayers": Table 1

| Drug | Pathway affected | MDCK WT | MDCK E-cadherin KO |
|---|---|---|---|
| No drug | -- | Extensile | Contractile |
| Blebbistatin (5 µM) | Non-muscle Myosin II | -- | Contractile |
| Blebbistatin (20 µM) | Non-muscle Myosin II | Extensile | Extensile |
| Y27632 (25 µM) | ROCK 1 and 2 | Extensile | Extensile |
