## Supplementary material for "Nature of active forces in tissues: how contractile cells can form extensile monolayers": Methods

This PDF contains:
Methods
Supplementary Figures 1-11
Table1
Video legends

### Methods

**Cell culture and reagents** MDCK WT (ATCC CCL-34) cells, MCF7A cells (ATCC HTB-22),MDCK overexpressing LifeAct Ruby, MDCK E-cadherin Knock-Out (KO) cells and shMCF7A E-cadherin KD cells were cultured in DMEM (containing Glutamax, High Glucose, and Pyruvate, Life Technologies) supplemented with 10% foetal bovine serum (Life Technologies) and 1% penicillin-streptomycin (Life Technologies) at 37°C with 5% $CO_2$. For cell migration experiments, cells were left to spread overnight before imaging so that the cells form a complete monolayer. Prior to imaging, normal culture media (DMEM) was changed to low glucose DMEM (containing Pyruvate, Life Technologies) in order to minimize cell division as cell divisions were known to generate extensile flow.[1] In our coculture mixing experiments, in order to ensure we have a mixed population at the start of imaigng, cells were plated with low Ca2+ media (no FBS) for 3 hours, and changed to normal media once the cells attach. For immunofluorescent stainings, cells were fixed with 4% paraformaldehyde (PFA), permeabilized with 0.5% Triton-X 100 for 5 minutes, blocked with 1% BSA/PBS for 1 hour, and incubated with primary antibody overnight at 4°C. The samples were then incubated with secondary antibody and Hoescht (Thermo Fisher)(1:10000) for 1 hour and mounted on Mowiol 4-88 (Sigma Aldrich C2081) before imaging. The primary antibodies used were directed against E-cadherin (24E10- Cell Signaling Technology; DECMA1- Sigma Aldrich) (1:100), cadherin 6(1:50),[2] paxillin (Y133- Abcam) (1:100), pMLC2 (Cell Signaling) (1:100), vinculin (kindly provided by Marina Glukhova) (1:2) ,[3] $\alpha$-catenin (Sigma Aldrich)(1:100), $\beta$-catenin (BD Biosciences) (1:100), ZO1 (a generous gift from Sylvie Robin) (1:50), YAP (Santa Cruz Biotechnology) (1:100). Anti-mouse, anti-rat, and anti-rabbit secondary antibodies conjugated with Alexa (488 or 568)(used at 1:200 dilution), Alexa 647 (1:50) conjugated phalloidin were purchased from Life Technologies. A slightly different fixation protocol was used to stain vinculin at the cell-

cell junction and focal adhesion sites. In order to label vinculin at cell-cell contact sites, cells were fixed with a mix of 4% PFA and 0.5% Triton-X 100 for 1 minute 30 seconds, followed by fixation with 4% PFA for 10 minutes. While staining for vinculin at focal adhesion sites, cells were fixed with 4% PFA for 10 minutes, followed by premeabilization with 0.5% Triton-X 100 for 10minutes. For experiments requiring inhibition of contractility blebbistatin 5 μM, 20 μM (Sigma Aldrich) or Y27632 25 μM (Sigma Aldrich) were added just before imaging.

**Generation of E-cadherin KO cell line** MDCK E-cadherin KO stable cells were generated using a CRISPR-Cas9 double nickase plasmid (Santa Cruz Biotechnology). The following gRNA sequences were used: TGATGACACCCGATTCAAAG and ATAGGCTGTCCTAG-GTAGAC. Around 2 million cells were electroporated (Neon Transfection System Invitrogen) with 3μg of plasmid in one pulse of 20 ms and at 1650V. Twenty four hours later, cells were selected by adding 2.5 μg/ml puromycin in the culture media. Fourty eight hours later, GFP positive single cells were sorted in 96 well plates by flow cytometry using Influx 500 sorter-analyzer (BD BioSciences). The clonal populations were then selected based on the absence of E-cadherin by immunofluorescence staining.The absence of E-cadherin in the clones generated was confirmed by Western blot analysis of protein extracts (Extended Figure 2b).

**Generation of siRNA E-cadherin KD cells** Lipofectamine RNAiMAX (Invitrogen) was used for siRNA transfection. siRNA sequences were control (on-target plus nontargeting pool), UGGUUUACAUGUCGACUAA. siRNA against E-cadherin : GGGACAACGUUUAUUACUA was used. The levels of E-cadherin was confirmed by Western blot analysis of protein extracts (Extended Figure 3a).

**Live cell and fixed sample imaging** Live imaging was performed with a 10X objective on BioStation IM-Q (Nikon) at 37°C and 5% CO2. Images are acquired every 10 min. For migration experiments, just the phase contrast images were captured every 10 min. For TFM experiments, phase contrast and fluorescent beads were imaged.

**Calculation of cell area, aspect ratio and molecular markers** The cellular area and aspect ratio were obtained from time lapse imaging of phase contrast images. Cells were then segmented using MorphoLibJ,[4] an ImageJ plugin for cell segementation. The area and length of paxillin was obtained by fitting them with an ellipse. Nuclear-cytoplasmic ratio of YAP intensity was quantified using an in-house ImageJ script. If the nucleus-cytoplasmic ratio was greater than 1.1 then YAP was considered to be nuclear while a value less than 0.99 was considered to be more cytoplasmic while any value in between was considered to be uniformly distributed through the cell.

**Western Blot** Proteins for MDCK cells were extracted using RIPA buffer without SDS (50mM Tris pH 7.5, 150mM NaCl, 1% NP40, 5mM EDTA, 1mM Na3VO4, 10mM NaF, 1mM PMSF, 1X protease inhibitor cocktail (Roche) and 1X phosphatase inhibitor (Phosphostop, Roche). Proteins from MCF7 cells were extracted using sample buffer (50mM Tris pH 7.5, SDS 2%, Glycerol 10%, Bromophenol blue 0.1%, Dithiothreitol 400nM, sterile water). Protein concentration was quantified by a Bradford assay (BioRad). 30 μg of protein were loaded onto NuPage 4-12% Bis-Tris gel using a mini gel tank and dry transferred using iBlot transfer system (Invitrogen). Non-specific sites were blocked using 5% non fat dry milk in 0.1% PBS Tween. For MLC total, blots were blocked with BSA/TBST (Tris buffered saline with Tween 20). Primary antibodies were diluted in PBS Tween at E-cadherin (24E10- 1:1000 for MDCK cells) (Santa Cruz, SC7870- 1:200 for MCF7 cells), $\alpha$-catenin (1:1000), $\beta$-catenin (1:1000), vinculin (gift from Marina Glukhova, 1:500), GAPDH (Protein Tech Europe 60004-1 for MDCK cells and abcam-1:5000, ab181603 for MCF7 cells- 1:500), alpha-tubulin (1:5000) (Sigma T9026) overnight on a shaker at 4 °C. Anti-MLC (Cell Signaling) antibodies were diluted in TBST. The blots were then washed 3-4 times for 10 minutes each in PBS 0.1% Tween or TBST (for pMLC2 and MLC total antibodies). They were then incubated with either Gampox, HRP linked (Sigma Aldrich, Pierce or Santa Cruz) or Dylight 800 linked secondary antibodies (ThermoFisher Scientific) for 2 hours. The blots were then washed three times with PBS 0.1% Tween or TBST for 10 minutes each. The blots were then revealed using CHEMIDOC MP (BioRad) using Super West Femto (34095 Thermo Scientific) or chemiluminescence.

**Traction force microscopy** Soft silicone substrates were prepared as described previously.[5] CyA and CyB were mixed in the ratio 1:1 and directly poured on glass bottom Petri dishes (fluorodish) in order to obtain a 100 μm thick layer. The substrate was cured at room temperature overnight on a flat surface. To ensure complete curing, the samples were cured at 80°C for 1 hour the next day. The surface was silanized using a solution of 5% APTES diluted in absolute ethanol for 5 min. The substrate was then washed with absolute ethanol and dried at 80°C for 10 min. 200 nm carboxylated fluorescent beads (Invitrogen) were diluted in deionized water solution at 1:500 for 10 min, washed with deionized water and dried at 80°C for 10 min. We then coated these substrates with 50 μg/ml fibronectin for 1 hour and washed with PBS prior to cell seeding. Around 200,000 cells were seeded in each petridish 40-50 min and washed with media when enough cells have attached. The cells were let to attach and spread overnight. The cells are imaged for 24 hours and at the end of the experiment, cells were removed with the addition of 500 μL of 10% SDS in the media so that the resting position of beads can be obtained.

**Laser ablation** The ultraviolet laser ablation system(355nm, 300ps pulse duration, 1 kHz repetition rate, PowerChip PNV-0150-100, team photonics)[6] was used to perform these experiments. MDCK overexpressing Lifeact-Ruby was employed and the apical section of the cell with the highest junctional intensity was used for ablation. Junctions between two tricellular contacts were ablated using the following parameters: laser power - 120nW, exposure time -

0.3sec and imaging interval - 2.2sec. Recoil velocity was computed by i) calculating the internodal distance after ablation by tracking the cartesian coordinates of the tricellular junctional nodes using MTrackJ plugin in Fiji [7] ii) fitting the calculated internodal distance into a single/double exponential function iii) obtaining recoil velocity using derivative of the function through a custom-made MATLAB algorithm.[6] The length of the junction was measured by drawing a line ROI in Fiji.[8] Junctional length and associated recoil velocity were plotted.

**Soft polyacrylamide gel patterning** Glass coverslips were plasma activated and coated with 0.1mg/ml PLL-g-PEG (SuSoS Technology). 1mm diameter circles were patterned on the passivated glass coverslips using deep UV and incubated the glass coverslips with 20 μg/ml fibronectin for 30 minutes. After incubation, glass coverslips were rinsed in 1x PBS to remove excess protein. Simultaneously, silanization of another set of glass coverslips were performed by plasma activation of clean coverslips followed by incubation with an ethanol solution containing 2% (v/v) 3-(trimethoxysilyl) propyl methacrylate (Sigma-Aldrich, St Louis, Missouri, USA) and 1% (v/v) acetic acid. The silanised coverslips were heated at 120°C. Freshly made polyacrylamide (PA) mix (7.5% acrylamide, 0.075% bis-acrylamide, 0.05% ammonium persulphate and 0.75 μl TEMED) was sandwiched between the patterned glass coverslip and silanized coverslip. The acrylamide, bis-acrylamide concentration was the same as,[9] to generate 2.3 kPa PA gels. After polymerization, the patterned coverslips were peeled off to reveal the patterns of protein on PA gels. Samples were kept submerged in 1x PBS until cell seeding.

**Analysis methods** Nematic analysis: Orientation field and defects were detected as described previously.[10] In short, the largest eigenvector of the structure tensor was obtained for each pixel while the orientation of cells were obtained using a plugin on ImageJ called OrientationJ. Using the winding number parameter, we identify defects within the monolayer. Then we obtain the local nematic order parameter tensor Q (which is averaged over a region of 3-4 cells). The largest eigenvector of Q was taken to be the orientation of 3-5 cells and plotted as red lines over the phase image to ensure that orientation identified is correct. Using this Q value automated defect detection can be done using the winding number parameter thereby detecting the various defects (+1/2, -1/2, +1 and -1) although we have more +1/2 or -1/2 defects. In order to reduce noise, only stable defects which are found in at least six consecutive frames (60 mins) are used in the following analysis as described in .[10] In addition, we manually tracked a few defects over time to verift their movement direction.

Velocity analysis: We use PIVlab (a tool implemented using Matlab) to analyse the velocity of cellular monolayers. An interrogation window of 64x64 ($40.96\,\mu m$) and 32x32 pixels ($20.48\,\mu m$) with an overlap of 50% were used for this analysis. Outlier vectors were manually removed and a local standard deviation filter was applied. The velocity correlation length was obtained using the formula as detailed in.[11]

Strain rate and stress measurement: Having identified the location of defects, we obtain the velocity field around the defects identified by aligning these defects. The strain rate was calculated from the gradient of the velocity field as $\dot{\varepsilon} = \vec{\nabla}\vec{v}$. By plotting the strain rate and velocity around the defect, we can characterize the system as an extensile or contractile system. For force measurement, the beads images obtained during cell migration are merged with the reference bead images obtained after SDS treatment. The images are stabilized using the Image Stabilizer plugin in ImageJ after which the illumination is corrected to remove background noise. We then obtain the displacement of beads using PIV of interrogation window 32x32 pixel with an overlap of 50%. Using the ImageJ plugin FTTC [12] we correlate the bead displacement to traction forces using a regularization parameter of $9 \times 10^{-9}$. Stress within the monolayer was estimated using Bayesian Inversion Stress Microscopy (BISM) with a regularisation parameter of $\Lambda = 10^{-6}$ .[13] This method obtains the stress directly from traction forces irrespective of epithelial rheology.

Isotropic stress was obtained as half the trace of the stress tensor $((s_{xx} + s_{yy})/2)$ in the tissue. Since the stress values obtained through this method are not reliable very close to the boundary, only defects in the center of the monoloyer have been taken into account in these calculations. The heatmaps obtained for strain rate and stress were smoothed through linear interpolation.

**Statistics** Differences between data were assessed using unpaired t-test implemented in Matlab and further verified using Graphpad Prism. On the plots, n.s.: not significant, $*p < 0.05$, $**p < 0.01$, $***p < 0.001$ and $****p < 0.0001$. Pearson coefficient was calculated using GraphPad Prism.

**Computational model** The model used in this manuscript is the extension of a recently developed phase-field model that has been shown to reproduce active nematic behavior in cellular tissues[14] and has been quantitatively compared with experiments showing coherent oscillations in confined epithelial monolayers.[15] We consider a two-dimensional tissue and describe each cell $i$ independently by a phase-field $\phi_i$, where $\phi_i \simeq 1$ indicates the interior of the cell and $\phi_i \simeq 0$ its exterior. The interface of each cell thus lies at $\phi_i = 1/2$. The phase-field dynamics is given by a Cahn-Allen type evolution equation:

$$\partial_t \phi_i + \vec{v}_i \cdot \vec{\nabla} \phi_i = -\frac{\delta \mathcal{F}}{\delta \phi_i}, \tag{1}$$

where $\vec{v}_i$ is the cell velocity that is determined from an over-damped equation for force balance as detailed below. $\mathcal{F}$ is the free energy that determines both mechanical properties of the cell - including cell stiffness and compressibility - and details of the passive interactions between the cells. As such the free energy $\mathcal{F} = \mathcal{F}_{\text{G-L}} + \mathcal{F}_{\text{area}} + \mathcal{F}_{\text{rep}} + \mathcal{F}_{\text{adh}}$ is composed of (i) Ginsburg-Landau term $\mathcal{F}_{\text{G-L}}$, that stabilises the interface, (ii) a soft constraint for area conservation $\mathcal{F}_{\text{area}}$, that penalises deviations from an initial circular morphology of the cell, (iii) $\mathcal{F}_{\text{rep}}$, which prevents two phase-fields from overlapping, and $\mathcal{F}_{\text{adh}}$, which increases the contact line between neighboring cells:

$$F_{\text{G-L}} = \sum_i \int \mathrm{d}\vec{x}\, \gamma \left\{ \frac{30}{\lambda^2}\, \phi_i^2 (1-\phi_i)^2 + (\vec{\nabla}\phi_i)^2 \right\}, \tag{2}$$

$$F_{\text{area}} = \sum_i \frac{\mu}{\pi R^2} \left( \pi R^2 - \int \mathrm{d}\vec{x}\, \phi_i^2 \right)^2, \tag{3}$$

$$F_{\text{rep}} = \sum_i \sum_{j \neq i} \frac{30\kappa}{\lambda^2} \int \mathrm{d}\vec{x}\, \phi_i^2 \phi_j^2, \tag{4}$$

$$F_{\text{adh}} = \sum_i \sum_{j \neq i} \frac{30\omega}{\lambda^2} \int \mathrm{d}\vec{x}\, \vec{\nabla}\phi_i \cdot \vec{\nabla}\phi_j, \tag{5}$$

where $\lambda$ sets the interface width, $\gamma$ sets the stiffness, $\mu$ determines cells compressibility, and $\kappa$, $\omega$ set the strength of repulsion and adhesion between two phase-fields, respectively. For the details of these free energy definitions, the reader is referred to recent reviews of phase-field models[16,17] and to[14,15,18,19] for recent implementations. Note that because here we model highly-packed, confluent tissues we do not introduce any thermodynamic attraction between the cells.

**Force balance.** We consider over-damped dynamics of cells moving on a substrate:

$$\xi \vec{v}_i = \vec{F}_i^{\text{tot}}, \tag{6}$$

where $\xi$ is the friction coefficient between the cells and the substrate, and $\vec{F}_i^{\text{tot}}$ denotes the total forces acting on each cell. This encompasses self-propulsion forces generated by the cell $\vec{F}_i^{\text{sp}}$ and the interaction forces $\vec{F}_i^{\text{int}}$ that a cell experiences from the neighbouring cells in the monolayer.

The self-propulsion force of an individual cell is achieved through an intricate coordination of actin polymerisation and myosin contractility. First, actin polymerisation at the cell front results in the formation of (lamelipodium) protrusions that deform the cell. Myosin contractility then retracts the cell rear to propel the cell forward. To account for the protrusion effects we introduce a polarity force $\alpha\vec{p}_i$, that is distributed over the front edge of the cell in the direction of the cell polarity $\vec{p}_i$, where $\alpha$ sets the strength of the polarity force. To account for the cell contractility, we then introduce a contractile stress $\zeta_Q \mathbf{Q}_i$, where $\zeta_Q$ is the strength of the contractility and $\mathbf{Q}_i = \vec{p}_i^{\mathrm{T}}\vec{p}_i - \frac{\mathbb{I}}{2}\vec{p}_i^2$ is the tensor that characterises the orientation of the polarity: the largest eigenvector of $\mathbf{Q}_i$ is $\vec{p}_i$ meaning that the contractile stress acts along the direction of protrusions formation. Considering that the vectors $\vec{\nabla}\phi_i$ describe the normal vector to the interface we obtain the following expression for the self-propulsion force:

$$\vec{F}_i^{\mathrm{sp}} = \alpha\vec{p}_i + \int \mathrm{d}\vec{x} \Big( - \zeta_Q \phi_i \mathbf{Q}_i \Big) \cdot \vec{\nabla}\phi_i, \tag{7}$$

where matrix multiplication is implied in the last term.

Next we consider the interaction stresses $\boldsymbol{\sigma}_i^{\mathrm{int}}$ to define the interaction forces $\vec{F}_i^{\mathrm{int}} = \int \mathrm{d}\vec{x}\, \phi_i \vec{\nabla} \cdot \boldsymbol{\sigma}^{\mathrm{int}} = -\int \mathrm{d}\vec{x}\, \boldsymbol{\sigma}^{\mathrm{int}} \cdot \vec{\nabla}\phi_i$. Note that $\vec{\nabla}\phi$ is only non-zero at the interfaces between the cells and as such the interaction force is acting at the cell-cell interfaces. We decompose the interaction stress in between the cells into passive and active contributions $\boldsymbol{\sigma}_i^{\mathrm{int}} = \boldsymbol{\sigma}_i^{\mathrm{passive}} + \boldsymbol{\sigma}_i^{\mathrm{active}}$: the passive contribution has a thermodynamic nature and is calculated from the free-energy:

$$\boldsymbol{\sigma}_i^{\mathrm{passive}} = \sum_i -\frac{\delta \mathcal{F}}{\delta \phi_i} \tag{8}$$

where $\delta$ denotes total derivative. The active contribution leads to the force generation between cells at their interface through adherens junction. Following our recent work[14] this takes the form

$$\boldsymbol{\sigma}_i^{\mathrm{active}} = -\zeta_S \sum_j \phi_j \mathbf{S}_j, \tag{9}$$

where $\mathbf{S}_i = -\int \mathrm{d}\vec{x}\, (\vec{\nabla}\phi_i)^{\mathrm{T}} \vec{\nabla}\phi_i$ is the deformation tensor for cell $i$, characterising the anisotropy of the cell shape such that the largest eigenvector of $\mathbf{S}_i$ corresponds to the direction of the elongation of the cell.

**Alignment dynamics.** We now introduce the dynamics of the cell polarity, modeling the mechanism that determines in which direction the polar force should act. There are many ways to introduce a dynamics of the polarisation.[20] One such way is through modeling the phenomenon of "contact inhibition of locomotion (CIL)", aligning the polarity of the cell to the direction of the total interaction force acting on the cell:[21] We define the dynamics of the polarisation to be given by

$$\partial_t \theta_i = -J|\vec{F}_i^{\text{int}}|\Delta\theta_i + D_r\,\eta, \tag{10}$$

where $\theta_i \in [-\pi, \pi]$ is the angle that the polarity vector is pointing at – such that $\vec{p}_i = (\cos\theta_i, \sin\theta_i)$ – and $\eta$ is a Gaussian white noise with zero mean, unit variance, and the rotational diffusivity $D_r$. The positive constant $J$ sets the time scale for the alignment of the polarity to the total interaction force, as was suggested theoretically[21] and has been recently confirmed in the experiments on epithelial cells.[15] As explained in[15] this model of alignment has the advantage that (i) it contains an explicit timescale and (ii) does not require that a cell 'knows' about the position of its neighbours.

**Simulation details** We simulated equation (1) using a finite difference scheme on a square lattice with a predictor-corrector step. Throughout this article, we used the following numerical values for the simulation parameters: $R = 8$, $\lambda = 3.0$, $\gamma = 0.04$, $\mu = 4.0$, $\kappa = 0.4$, $\omega = 0.0$ $\xi = 1$, $\alpha = 0.15$, $\zeta_Q = -0.08$, and $\zeta_S = 0.02$ for the wild type case, while $\zeta_S = 0.0$ for the case with no cell-cell interaction stresses. We simulated square domains of edge length $W = 100, 200, 400$ lattice sites with a packing fraction of $\Phi = 1.2$ cells and set $D_r = 1 \times 10^{-3}$ and $J = 0.05$.
